# Joint sequence & chromatin neural networks characterize the differential abilities of Forkhead transcription factors to engage inaccessible chromatin

**DOI:** 10.1101/2023.10.06.561228

**Authors:** Sonny Arora, Jianyu Yang, Tomohiko Akiyama, Daniela Q. James, Alexis Morrissey, Thomas R. Blanda, Nitika Badjatia, William K.M. Lai, Minoru S.H. Ko, B. Franklin Pugh, Shaun Mahony

**Author notes:** Equal contributions (Sonny Arora, Jianyu Yang).

## Abstract

The DNA-binding activities of transcription factors (TFs) are influenced by both intrinsic sequence preferences and extrinsic interactions with cell-specific chromatin landscapes and other regulatory proteins. Disentangling the roles of these binding determinants remains challenging. For example, the FoxA subfamily of Forkhead domain (Fox) TFs are known pioneer factors that can bind to relatively inaccessible sites during development. Yet FoxA TF binding also varies across cell types, pointing to a combination of intrinsic and extrinsic forces guiding their binding. While other Forkhead domain TFs are often assumed to have pioneering abilities, how sequence and chromatin features influence the binding of related Fox TFs has not been systematically characterized.

Here, we present a principled approach to compare the relative contributions of intrinsic DNA sequence preference and cell-specific chromatin environments to a TF’s DNA-binding activities. We apply our approach to investigate how a selection of Fox TFs (FoxA1, FoxC1, FoxG1, FoxL2, and FoxP3) vary in their binding specificity. We over-express the selected Fox TFs in mouse embryonic stem cells, which offer a platform to contrast each TF’s binding activity within the same preexisting chromatin background. By applying a convolutional neural network to interpret the Fox TF binding patterns, we evaluate how sequence and preexisting chromatin features jointly contribute to induced TF binding.

We demonstrate that Fox TFs bind different DNA targets, and drive differential gene expression patterns, even when induced in identical chromatin settings. Despite the association between Forkhead domains and pioneering activities, the selected Fox TFs display a wide range of affinities for preexiting chromatin states. Using sequence and chromatin feature attribution techniques to interpret the neural network predictions, we show that differential sequence preferences combined with differential abilities to engage relatively inaccessible chromatin together explain Fox TF binding patterns at individual sites and genome-wide.

## INTRODUCTION

Pioneer transcription factors have an ability to bind DNA targets in relatively inaccessible chromatin, thereby establishing competency for the binding of other transcription factors during development [1–4]. But pioneer factors are also known to bind to different sites in different cell types [5], suggesting a dependency on cell-specific regulatory environments and blurring the lines between pioneer and non-pioneer TFs. Understanding where TFs lie along the spectrum of pioneering abilities will require characterization of how TFs’ DNA-binding patterns are influenced by both their intrinsic sequence preference and their interactions with cell-specific chromatin environments and other regulators [6–8]. However, we currently lack principled approaches for measuring and comparing the sequence and chromatin determinants of TF binding specificity across TFs.

Amongst the earliest recognized pioneer factors were the Forkhead domain-containing FoxA family of TFs. FoxA1 and FoxA2 interact with nucleosomal binding sites during early endodermal development, enabling other transcription factors to bind to lineage-specific enhancer sequences [9–12]. The Forkhead domain has a similar structural fold to the H1 linker histone [13], leading to suggestions that Fox TFs compete with and displace H1 histones at target sites [14]. Most Fox DNA contacts occur through an α-helix and two flanking loops that are positioned on a single side of the DNA, making the Forkhead domain compatible with binding to nucleosome-wrapped DNA [15].

Despite their demonstrated abilities to engage nucleosomal targets, FoxA TFs bind to extensively different sets of sites in different cell types and at different points in development [5,16–20]. Several studies have tried to explain this apparent paradox by proposing that FoxA binding sites are pre-marked or potentiated by lineage-specific regulatory events. Cell-specific FoxA binding activities are associated with a variety of preexisting chromatin states, including preexisting DNA hypomethylation [18], active histone modifications [16,20], prior binding of other TFs (including other Forkhead paralogs) [21,22], and cooperative interactions with other co-expressed TFs [19,23–25]. For example, our previous analysis of FoxA1 ChIP-exo crosslinking patterns in estrogen-treated MCF7 cells suggests that a subset of FoxA1 binding sites occur indirectly via interactions with Estrogen Receptor alpha in that cell type [26]. Even when endogenously co-expressed in adult livers, the closely related paralogs FoxA1 and FoxA2 bind to different sites and regulate different genes, possibly via alternate associations with other regulators [27]. Overall, a wealth of data suggest that FoxA pioneering activities are not absolute, and instead cell-specific FoxA TF DNA-binding activities are determined by a mixture of intrinsic pioneering abilities and extrinsic cell-specific influences in the chromatin environment.

Outside of the FoxA subfamily, relatively little is known about how other Forkhead domain TFs determine their binding sites. There are over 40 Fox TF paralogs encoded in mammalian genomes, many of which are documented to play crucial regulatory roles in development and disease across diverse tissue types [28,29] (**Fig. S1A**). All Fox TFs share similar Forkhead DNA-binding domains with similar predicted structural folds. Thus, all Fox TFs might be expected to share the same structural properties that are hypothesized to enable FoxA TFs to engage nucleosomal DNA. Indeed, some Fox paralogs such as FoxE1 and FoxO1 bind individual nucleosomal sites and actively decondense chromatin [30,31]. However, at least one Fox paralog – FoxP3 – is not a pioneer factor during T cell development, instead being highly influenced by the preexisting binding of other regulators [32]. The conservation amongst Forkhead domains is also reflected in their DNA-binding preferences; most mammalian Fox TFs favor sequences containing a TGTTTA motif *in vitro* [33,34]. Interestingly, some Fox paralogs, and even some splice isoforms, can bind drastically different DNA-binding motifs in cellular contexts [35,36]. Therefore, it is generally unclear whether Fox TFs should be expected to bind similar sequences and share similar pioneering activities when expressed in the same cellular setting.

Here, we compare the abilities of several mammalian Fox TFs to engage their DNA-binding targets when introduced into a common chromatin environment. We focus on five Fox paralogs (FoxA1, FoxC1, FoxG1, FoxL2, and FoxP3) from across the phylogeny, each of which have similar Forkhead domain sequences and share high levels of identity at DNA-contacting residues (**Fig. S1B-C**). Importantly, all five selected Fox TFs have similar DNA-binding preferences according to DNA-binding experiments stored in the cis-bp database [34] (**Fig. 1A**). However, the five TFs share no conserved domains outside of the DNA-binding domain (**Fig. S1D**), and all five play distinct roles in development [28]. Therefore, despite sharing similar *in vitro* binding preferences, it is not clear whether the five selected Fox TFs would bind to similar sets of targets in a given cellular environment. To compare the DNA-binding activities of these selected Fox TFs, we first over-express each Fox TF in mouse embryonic stem (mES) cells, which do not endogenously express any of the selected factors (**Fig. 1B**). We then capture the high-resolution binding activities of each Fox TF using ChIP-exo and the regulatory outcomes using RNA-seq and ATAC-seq. The advantage of ectopically inducing their expression in a well-characterized chromatin environment is that it enables us to assess how DNA-binding activities relate to a common preexisting chromatin landscape.

**Figure 1:**
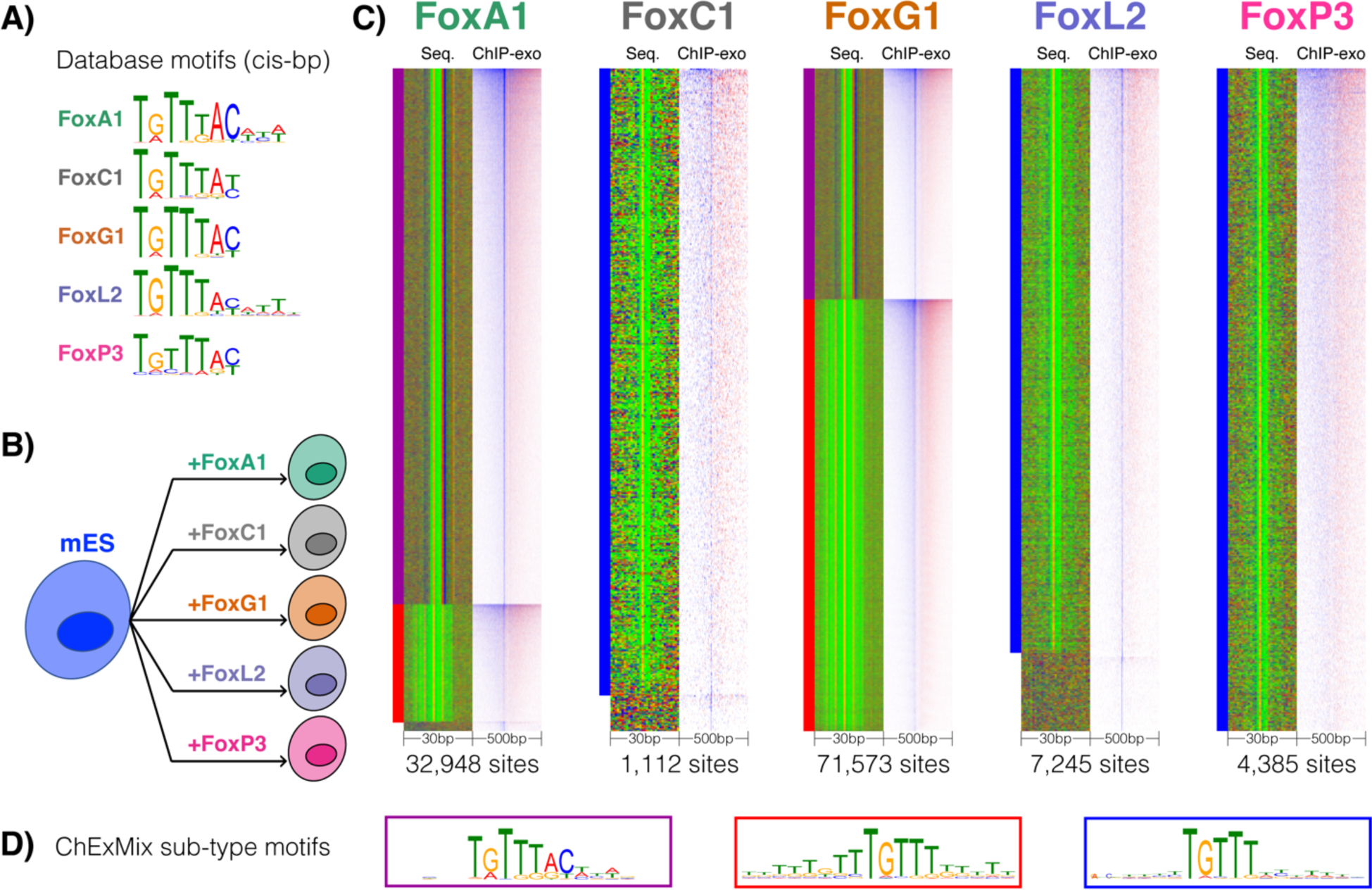
Fox TFs bind directly to cognate motifs when expressed in mES cells. **A)** Previously characterized *in vitro* DNA-binding preferences (sourced from cis-bp [34]) for each examined Fox TF. **B)** Expression of each selected Fox TF is induced in mES cells for 24hrs before ChIP-exo experiments. **C)** Heatmaps corresponding to ChExMix-defined Fox TF binding sites. For each Fox TF, two heatmap representations are shown. The heatmap on the right (“ChIP-exo”) displays the 5’ ends of ChIP-exo reads (blue = positive strand, red = negative strand) in 500bp windows centered on the ChExMix peak. The heatmap on the left shows DNA bases (red = A, blue = C, orange = G, green = T) in 30bp windows centered on the ChExMix peak. The ChExMix peaks for each Fox TF are clustered according to DNA-binding “subtypes” that are automatically detected by ChExMix based on the ChIP-exo read distributions and detected motifs. The purple, red, and blue bars to the left of the heatmap represents which type of Fox motif was detected in each ChExMix subtype, where the actual motifs corresponding to each subtype are shown in panel **D)**. Figure S3 shows the ChIP-exo profiles corresponding to each ChExMix subtype.

To model how Fox TF binding activities depend on sequence preferences and the preexisting mES chromatin environment, we developed a Convolutional Neural Network (CNN) that jointly models DNA and chromatin features to predict TF binding signals. Advancing on our previous approach for integrating sequence and preexisting chromatin predictors of TF binding [37], our CNN uses early integration of input features to predict the quantitative ChIP enrichment signal at binding sites. Using neural network feature attribution techniques on the trained CNN models, our approach can highlight and interpret how sequence and chromatin features together specify binding signals, both globally and at individual binding sites.

Our results demonstrate that, despite having similar *in vitro* DNA-binding preferences, Fox TFs display highly divergent DNA binding activities in a common mES chromatin environment. Using our CNNs, we show that differential Fox DNA-binding activities are only partially explained by DNA-binding motif differences in this cell type. Rather, the examined Fox TFs vary widely in their abilities to engage different chromatin landscapes.

## RESULTS

### Fox paralogs bind to different sites when expressed in mES cells

To capture the regulatory activities of Fox paralogs in a common chromatin environment, we took advantage of a collection of mES cell lines that contain inducible FLAG-tagged TFs [38–40]. We focus on five Fox TFs – FoxA1, FoxC1, FoxG1, FoxL2, and FoxP3 – which represent various points in the phylogeny of vertebrate Forkhead DNA-binding domains (**Fig. S1A**) and none of which are endogenously expressed in mES cells. The experimental system is intended to address how these Fox TFs integrate into a common chromatin environment that they have not encountered before, which gets to the heart of pioneering activities rather than addressing the biological role of Fox TFs in development. Each TF was over-expressed in (undifferentiated) mES cells for 24 hours (**Fig. 1B**), a timepoint chosen to balance between the goal of capturing the earliest DNA-binding activities in mES chromatin and the ability to assay robust ChIP signals (see Methods, **Table S1**, **Fig. S2**). We then characterized TF-DNA binding activities using ChIP-exo [41], which not only captures TF-DNA binding activities at single-base resolution, but also enables the detection of alternative protein-DNA binding modes via variation in DNA crosslinking patterns [26].

We analyzed our binding data using ChExMix, which is designed to find distinct binding site subtypes (‘modes’) in ChIP-exo data based on the motifs and ChIP-exo crosslinking patterns present at binding sites [26]. ChExMix revealed that the selected Fox TFs primarily bind directly to cognate motif instances in mES cells (**Fig. 1C-D**, **Fig. S3**). In contrast to our observations of FoxA1 binding in MCF7 cells [26], we do not observe any ChExMix binding modes that are anchored by non-Forkhead domain motifs, indicating that Fox-interacting partners may not be present or active in mES cells. However, the various Fox TFs displayed differential preferences in their primary cognate motifs. While cis-bp database entries suggest that all five Fox TFs should bind to a TGTTTAC motif, only FoxA1 and FoxG1 display a ChExMix binding mode containing this motif (**Fig. 1C-D**). The other binding modes detected by ChExMix, including the binding modes that account for the majority of FoxC1, FoxG1, FoxL2, and FoxP3 binding sites, contain a simpler core TGTTT pattern in one or more copies (**Fig. 1C-D**).

Quantitative differential binding analysis confirms that the five selected Fox TFs have highly divergent genomic binding distributions when expressed in mES cells (**Fig. 2A**). For example, 96% of FoxL2’s binding sites display significantly greater ChIP enrichment for FoxL2 than FoxA1, whereas 43% of FoxA1’s sites display significantly greater ChIP enrichment for FoxA1 than FoxL2. Fox TF differential binding preferences are more clearly visualized by clustering ChIP enrichment patterns at the top 5,000 peaks from each TF (**Fig. 2B**). Only a small subset of sites displays similar ChIP enrichment across all five Fox TFs. In contrast, larger fractions of sites are preferentially bound by FoxA1, FoxL2, or both FoxG1 and FoxP3.

**Figure 2:**
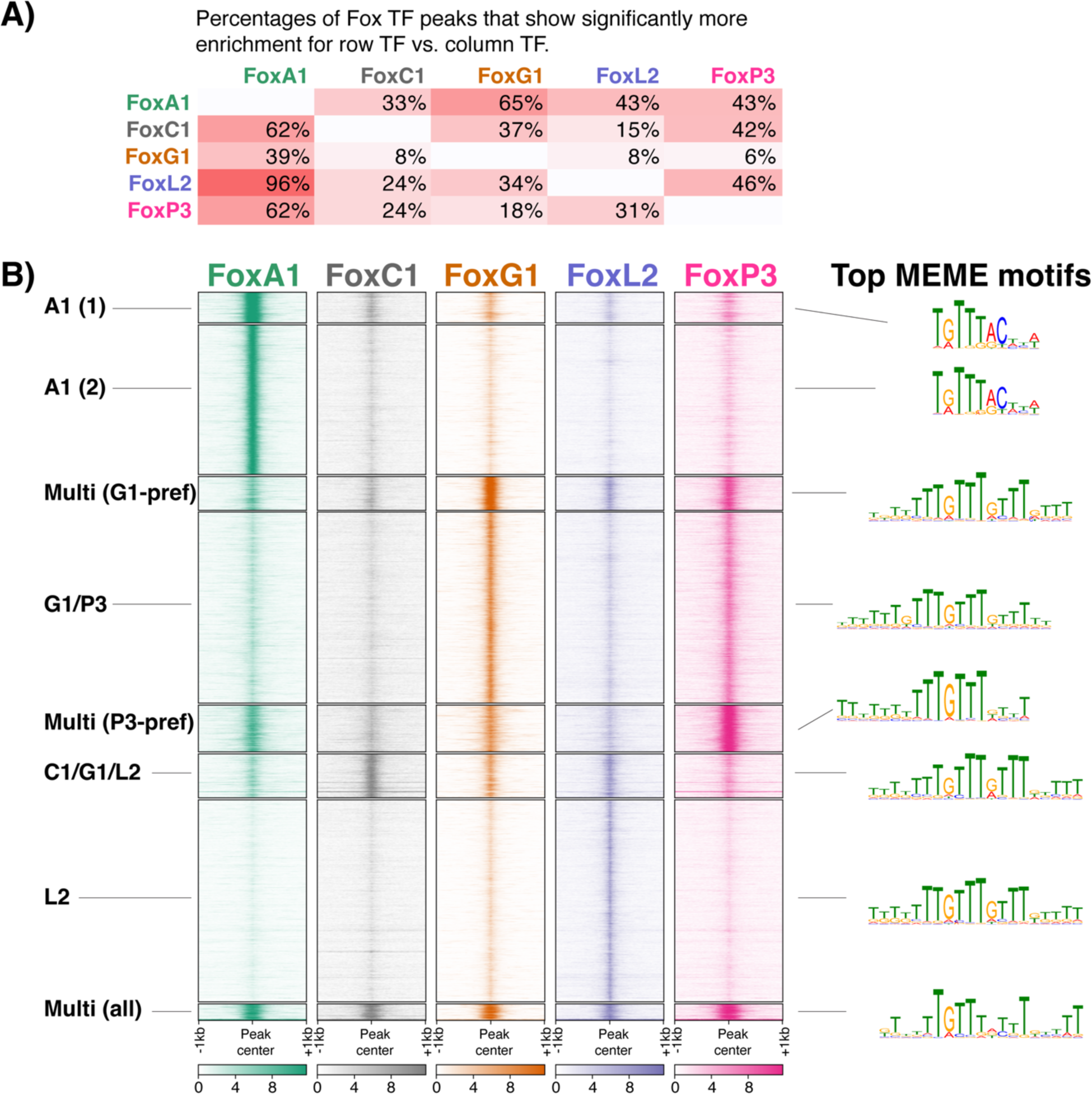
Fox TFs bind to distinct locations in mES cells. **A)** The table shows the percentages of the row TF’s top 5,000 peaks that display significantly greater enrichment in the row TF ChIP-exo experiment compared with the column TF experiment (as determined by edgeR differential binding analysis; fold>1.5, q<0.05). **B)** Unstranded ChIP-exo 5’ read enrichment plotted at clusters of Fox TF binding sites as determined by K-means clustering. The top-most significant motifs as discovered by MEME in each clusters’ binding sites are shown on the right. Figure S4 shows the top three MEME motifs for each cluster.

### Alternate sequence preferences do not fully explain Fox paralog binding differences

We investigated whether the differential binding patterns displayed by Fox TFs are explained by differential motif preferences. *De novo* motif-finding applied to each cluster of sites finds only limited variation in discovered patterns (**Fig. 2B**). The TGTTTAC pattern is the primary motif discovered in both clusters of sites that are preferentially bound by FoxA1, consistent with ChExMix analysis. However, the primary motif discovered in all other clusters is one containing a tandem-repeated TGTT pattern. *De novo* discovered secondary motifs do not explain the differences between binding site clusters either, with the exception of an Oct/Sox motif that is only discovered at a cluster of sites preferentially bound by FoxP3 (**Fig. S4**).

We next applied SeqUnwinder, a *k*-mer based multi-label discriminative motif-finder [42], to further elucidate sequence patterns that discriminate between Fox TF binding site clusters. SeqUnwinder finds several short sequence features that are preferentially enriched in different clusters (**Fig. S5A**). However, the discovered motifs are not strongly discriminative between clusters, both when examining the occurrence frequencies of each motif at sites in individual clusters (**Fig. S5B**) or when examining the performance of SeqUnwinder’s classification model in assigning held-out sites to each cluster (**Fig. S5C**). For example, the SeqUnwinder model does not find any features that strongly discriminate between the large cluster of sites that are preferred by FoxL2 and a smaller set of sites that are bound by FoxC1, FoxG1, and FoxL2.

Thus, while some Fox TFs (e.g., FoxA1) display preferential binding to specific cognate motif variants, sequence features alone do not fully explain the observed differential binding across Fox binding sites.

### Fox paralogs differentially associate with preexisting chromatin features

Our analyses of Fox TF binding site features suggest that the cognate DNA-binding motifs vary to some degree between Fox paralogs. However, the differential sequence features discovered at binding sites do not fully account for the differences in Fox TF binding patterns (**Fig. S5C**). We next asked whether chromatin features beyond the sequence level could help to explain the binding differences observed between Fox paralogs. All five Fox TFs are expressed in a common chromatin environment – that of mES cells – which has been extensively characterized by regulatory genomics assays. To profile how newly expressed Fox TFs associate with various preexisting chromatin features, we can therefore compare the various Fox TF binding sites to chromatin signals from the preexisting mES cells.

We first performed an overlap analysis between the top 2,000 peaks from each Fox TF and a selection of mES histone modifications and accessibility patterns (**Fig. 3A**). A majority of FoxP3 and FoxG1 sites overlap regions that already displayed enrichment for ATAC-seq or active histone modifications (H3K4me1/2, H3K27ac) in the preexisting mES cells. In contrast, relatively few FoxL2 or FoxC1 sites display overlap with these markers of preexisting regulatory activity. FoxA1 sites displayed an intermediate level of association.

**Figure 3:**
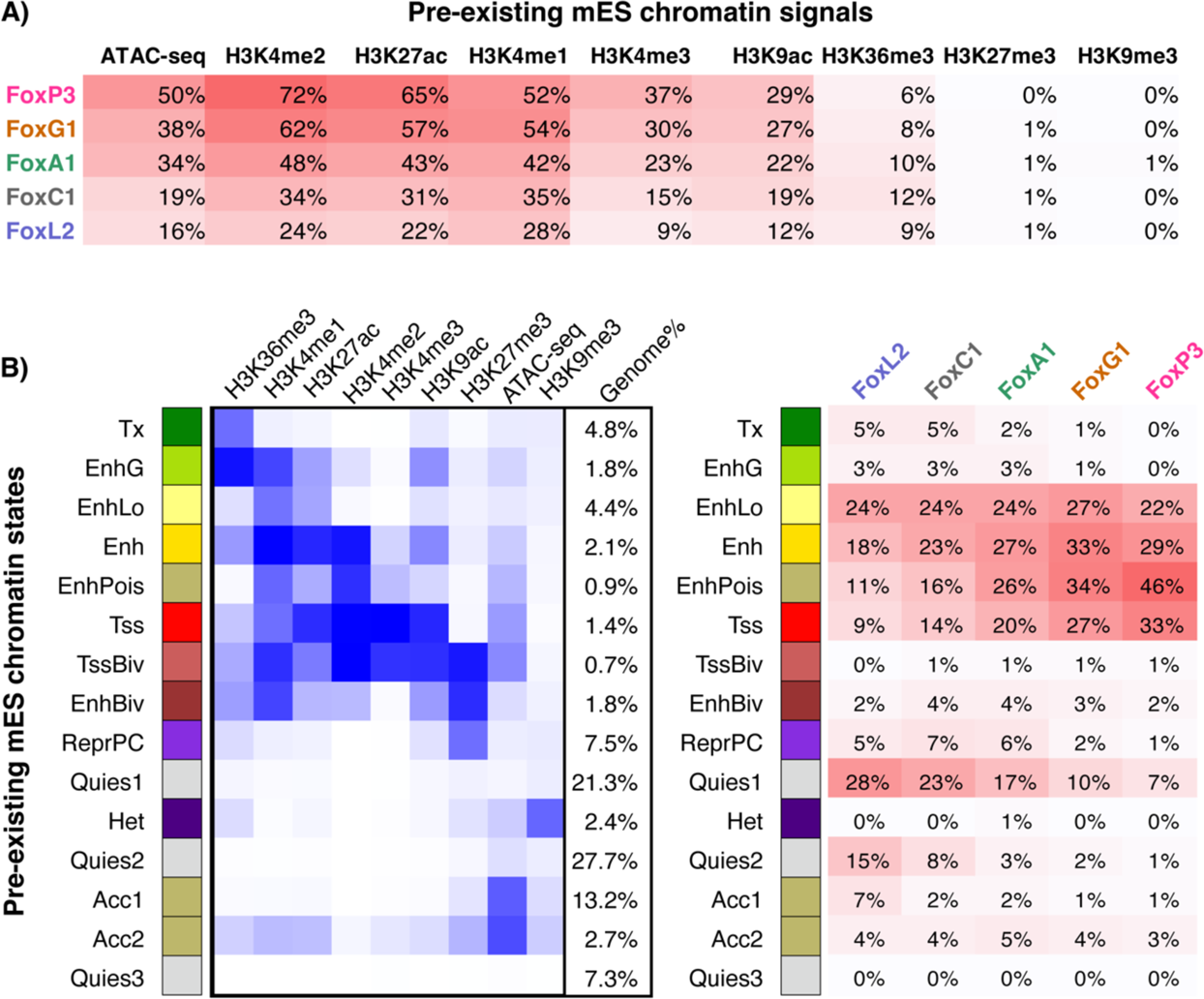
Fox TF binding sites differentially associate with preexisting mES chromatin features. **A)** Percentages of each Fox TF’s top 2,000 binding sites that are within 200bp of enriched domains for each listed mES chromatin feature. **B)** Left panel shows the chromatin states determined by ChromHMM using the mES chromatin features, where the blue shading shows the relative enrichment of each chromatin feature within each state. Right panel shows the percentages of each Fox TF’s top 2,000 binding sites that are within 200bp of regions covered by each state.

To explore associations with combinatorial chromatin patterns in more detail, we first characterized mES cell chromatin states using ChromHMM [43] and then calculated the percentages of Fox TF sites that are within 200bp of regions marked by each state (**Fig. 3B**). FoxP3 and FoxG1 display much higher levels of association with enhancer- and promoter-like states (“Enh”, “EnhPois”, “Tss”) compared with FoxL2 and FoxC1. Conversely, FoxL2 and FoxC1 display higher levels of association with quiescent regions (“Quies1” and “Quies2”) that are devoid of any included chromatin signals in mES cells. FoxA1 again displays intermediate levels of association with both active regulatory states and quiescent states. We note that these differential associations with preexisting enhancer- and promoter-like states are not merely due to some Fox TFs preferring to bind proximally to promoters; all five Fox TFs have similar distributions of distances to annotated TSSs (**Fig. S6A**) and similar associations with repeats (**Fig. S6B**).

We next looked for associations between Fox TF binding sites and the binding sites of other TFs that are already expressed in mES cells. While Fox TF binding sites overlap many mES TF binding sites at higher rates than random sites (**Fig. S7A**), the absolute percentage overlap rates are generally low (**Fig. S7B**). One exception to this trend is FoxD3, a Forkhead domain TF that is endogenously expressed in mES cells [44] (**Fig. S7B, Fig. S8**). Substantial fractions of FoxP3, FoxG1, and FoxA1 binding sites are pre-bound by FoxD3, consistent with a previous report that FoxD3 can potentiate FoxA1 binding sites in differentiation systems [21]. More generally, the associations between Fox TFs and mES TFs are consistent with the associations observed between Fox TFs and active chromatin features: FoxP3 and FoxG1 display generally higher associations with mES TF binding sites than FoxL2 and FoxC1, while FoxA1 shows intermediate association levels

In summary, our results demonstrate that FoxP3 and FoxG1 bind primarily to regions that were already accessible or displaying signatures of regulatory activity in the preexisting mES chromatin environment. FoxP3 and FoxG1 binding at these preexisting regulatory regions may be facilitated by non-specific interactions with prebound pluripotency TFs or FoxD3, consistent with the assisted loading model [45]. In contrast, FoxL2 and FoxC1 have a higher preference for quiescent regions that are devoid of regulatory activities in mES cells. While FoxA1 is a well-known pioneer TF, it displays intermediate levels of association with preexisting regulatory and quiescent regions.

### Representation of Fox TF binding activities is improved by accounting for the preexisting chromatin environment

Our results thus far led us to the hypothesis that the DNA-binding activities of some Fox paralogs, including FoxP3 and FoxG1, are more strongly influenced by the preexisting chromatin environment than others. Under this model, Fox TFs that are expressed in the same chromatin environment and that share similar DNA binding motifs could nevertheless bind to different sites based on differing abilities to bind to relatively inaccessible chromatin.

To further address this hypothesis, we developed a CNN architecture that predicts quantitative TF binding signal levels using DNA sequence and (optionally) chromatin features (**Fig. 4A**). We first trained CNNs to predict ChIP-exo read counts for each Fox TF from DNA sequence features alone. These “sequence-only” CNN models do not fully capture the dynamic range of the data (**Fig. 4B**). In particular, sequence-only models struggle to recapitulate the observed pattern of differential binding across the previously defined Fox binding site clusters (**Fig. 4C**, **Fig. S9**). For example, while FoxG1 displays a stronger binding preference for five of the eight clusters in the ChIP-exo data, this differential preference is not captured by the sequence-only FoxG1-trained CNN.

**Figure 4:**
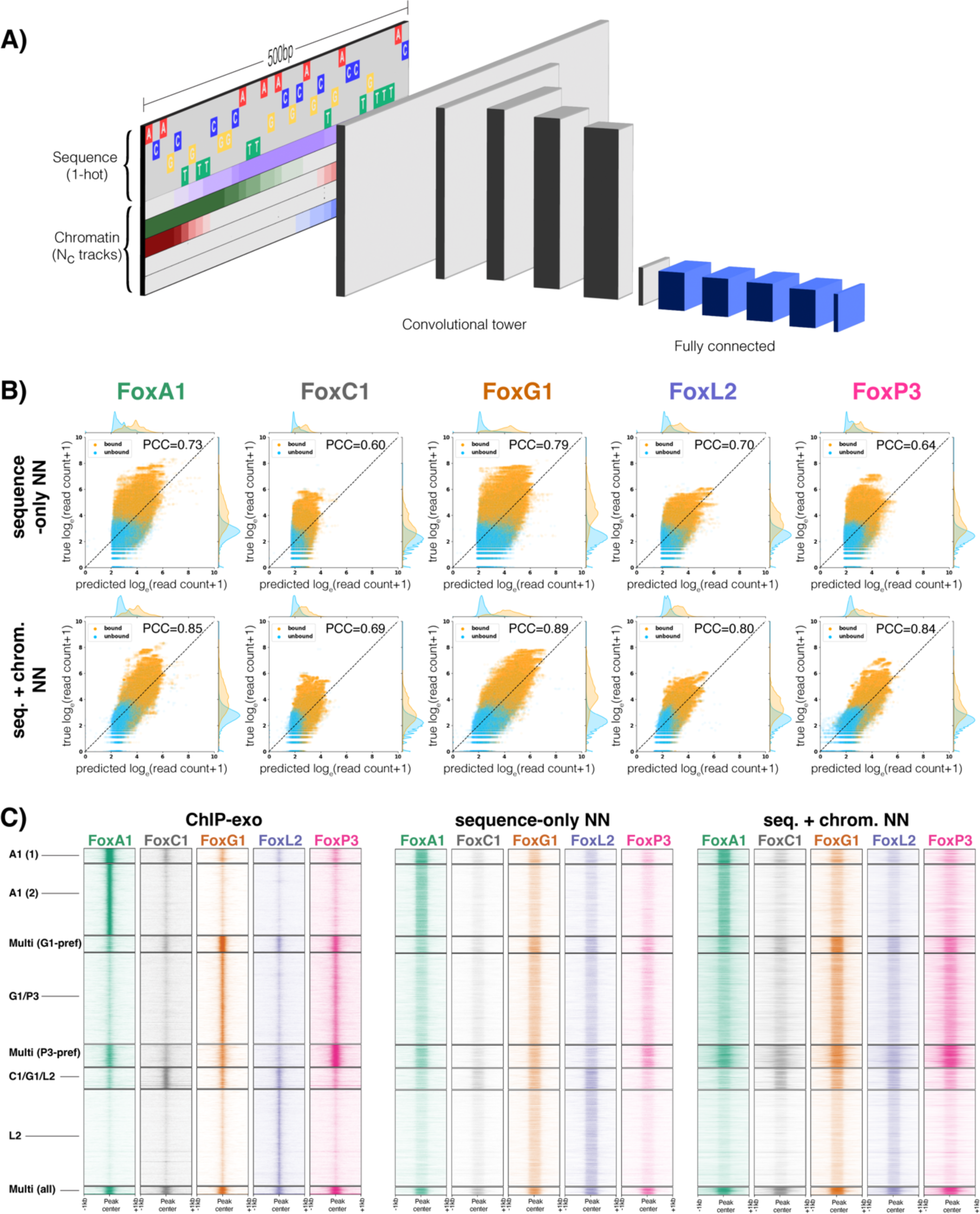
Convolutional neural networks trained with both sequence and chromatin features offer improved representation of Fox TF binding sites. **A)** Cartoon representation of the CNN architecture. **B)** Scatter plots showing bound (orange) and unbound (cyan) held-out test set sites for each Fox TF, where the y-axes represent the true log ChIP-exo read count, and the x-axes represent the predicted log read count from CNN models (trained with sequence only, or sequence and chromatin). **C)** Heatmap representations of CNN predicted read counts at the clusters of Fox TF binding sites displayed in Figure 2. Note that the CNNs predict read counts in 500bp windows, thus explaining the relative lack of resolution compared with the ChIP-exo data.

Next, to assess whether the preexisting mES chromatin environment is predictive of future induced Fox TF binding, we trained “sequence+chromatin” CNNs to predict ChIP-exo read counts using both sequence and mES chromatin features (ATAC-seq, H3K4me1, H3K4me3, H3K9me3, H3K27ac, H3K27me3, H3K36me3, H3K79me2, H3K122ac, high-concentration-MNase-seq, low-concentration-MNase-seq). Adding preexisting chromatin features to the CNN training process results in a substantial improvement in the ability to predict ChIP-exo read counts at held-out sites (**Fig. 4B**, **Fig. S10**). The ability to recapitulate differential binding across binding site clusters is also improved (**Fig. 4C**, **Fig. S9**). Improvements in ChIP-exo representation are particularly pronounced for FoxP3 and FoxG1, the factors whose binding sites are most highly associated with preexisting chromatin features. In contrast, mES chromatin features result in a smaller relative improvement in the representation of FoxL2 ChIP-exo read counts. These observations suggest that the preexisting chromatin environment is less predictive of FoxL2 binding than it is of FoxP3 and FoxG1 binding.

To further characterize how the CNNs integrate sequence and chromatin information, we used feature attribution approaches to examine how the various sources of information contribute to sequence+chromatin CNN predictions (**Fig. 5**). The FoxA1 CNN assigns higher contribution scores to DNA sequences at the center of peaks in the FoxA1-preferred clusters compared with sites in other clusters (**Fig. 5A**). As shown in **Fig. 2B**, the FoxA1-preferred clusters are associated with the presence of the TGTTTAC cognate motif. Thus, the FoxA1 CNN uses DNA sequence features to discriminate sites in FoxA1-preferred clusters from those in other clusters. However, our CNNs do not appear to find DNA sequence features that similarly discriminate between binding site clusters for the other Fox TFs. For example, all clusters appear to receive similarly positive DNA sequence contribution scores from the FoxG1 CNN, even though FoxG1 does not uniformly bind across all clusters (**Fig. 5A**). The TF-MoDISco package, which compiles sequence attribution patterns into interpretable motifs, finds similar motifs from the FoxC1, FoxG1, FoxL2, and FoxP3 CNN sequence attributions (**Fig. S11**). These observations are consistent with our conclusions from traditional motif analyses that sequence features alone do not explain differential Fox binding across the clusters (with the partial exception of FoxA1).

**Figure 5:**
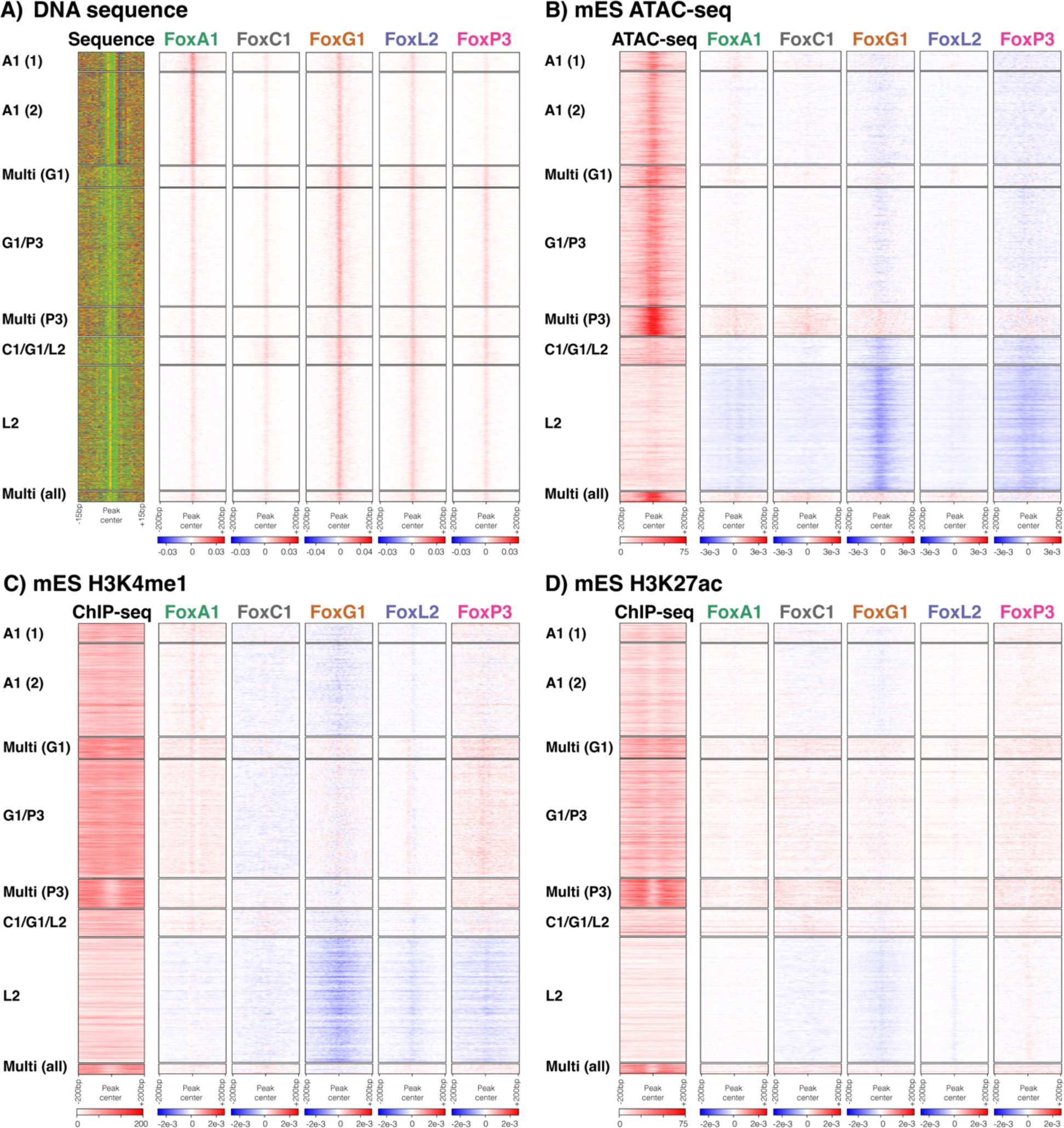
CNN feature attributions demonstrate the differential dependencies of Fox TFs on sequence and chromatin features. Each panel shows feature attribution results in 400bp windows centered on Fox TF binding sites for: **A)** sequence features; **B)** mES ATAC-seq chromatin features; **C)** mES H3K4me1 chromatin features; **D)** mES H3K27ac chromatin features. Sites are clustered as displayed in Figure 2. Positive feature attributions are shown in red, and negative in blue. To the right of each panel is a heatmap showing either the sequences in a 30bp window centered on the ChExMix peak or ATAC/ChIP-seq read depth in a 400bp window centered on the ChExMix peak. Figure S12 shows additional chromatin feature attribution heatmaps at the same sites.

In contrast to DNA sequence attribution scores, chromatin features appear to enable the CNNs to discriminate between binding site clusters in a more nuanced way. For example, the FoxG1 and FoxP3 CNNs interpret the low levels of mES ATAC-seq and H3K4me1 signals in the FoxL2-preferred cluster as a strongly negative feature that prevents the prediction of FoxG1 or FoxP3 binding sites at this cluster (**Fig. 5B**, **Fig. 5C**). Conversely, high levels of mES H3K4me1 or H3K27ac in clusters bound by FoxP3 are recognized as positive features by the FoxP3 CNN (**Fig. 5C**, **Fig. 5D**). Additional histone marks and MNase-seq signals also create chromatin attribution scores that distinguish between the Fox TF binding site clusters (**Fig. S12**). Thus, the FoxG1 and FoxP3 CNNs appear to be able to accurately predict binding at FoxG1- and FoxP3-preferred sites by modulating the high sequence scores seen across all Fox binding site clusters with chromatin scores that prefer permissive preexisting chromatin features.

How these general sequence and chromatin preferences contribute to Fox TF binding predictions at individual sites is illustrated using three examples (**Fig. 6**). **Fig. 6A** shows a FoxA1 binding site from the FoxA1-preferred cluster (chr3:103783444), where the FoxA1 CNN gives a high sequence attribution score to a TGTTTAC sequence in the center of the site. The FoxG1 and FoxL2 CNNs do not give a high score to the same sequence, suggesting that the presence of the specific TGTTTAC motif explains why the site is bound by FoxA1 and not the other two TFs. Similarly, **Fig. 6B** shows a FoxG1 binding site from the FoxG1-preferred cluster (chr8:34215670), where the FoxG1 CNN provides both positive sequence attribution scores to a TGTTT motif instance and a positive chromatin attribution score to overlapping mES ATAC-seq chromatin signals. The lack of positive attribution scores from the FoxA1 and FoxL2 CNNs suggest that the combination of sequence and accessibility features is favorable for FoxG1 but not FoxA1 or FoxL2 at this particular site. Finally, **Fig. 6C** shows a FoxL2 binding site from the FoxL2-preferred cluster (chr2:79050928), where the FoxL2 CNN provides positive sequence attribution scores to TGTTT instances in the region. The FoxA1 CNN does not score the same sequences positively, again pointing to FoxA1’s differential binding preference. However, the FoxG1 CNN provides high scores to the same TGTTT sequences, suggesting that it would predict this site as FoxG1-bound given sequence features alone. But these positive sequence scores are counter-balanced by negative chromatin attribution scores caused by the lack of mES ATAC-seq or H3K4me1 signals in this region. Thus, our joint sequence and chromatin CNNs can elucidate the specific contributions of sequence and chromatin features to explain differential binding activities at individual sites.

**Figure 6:**
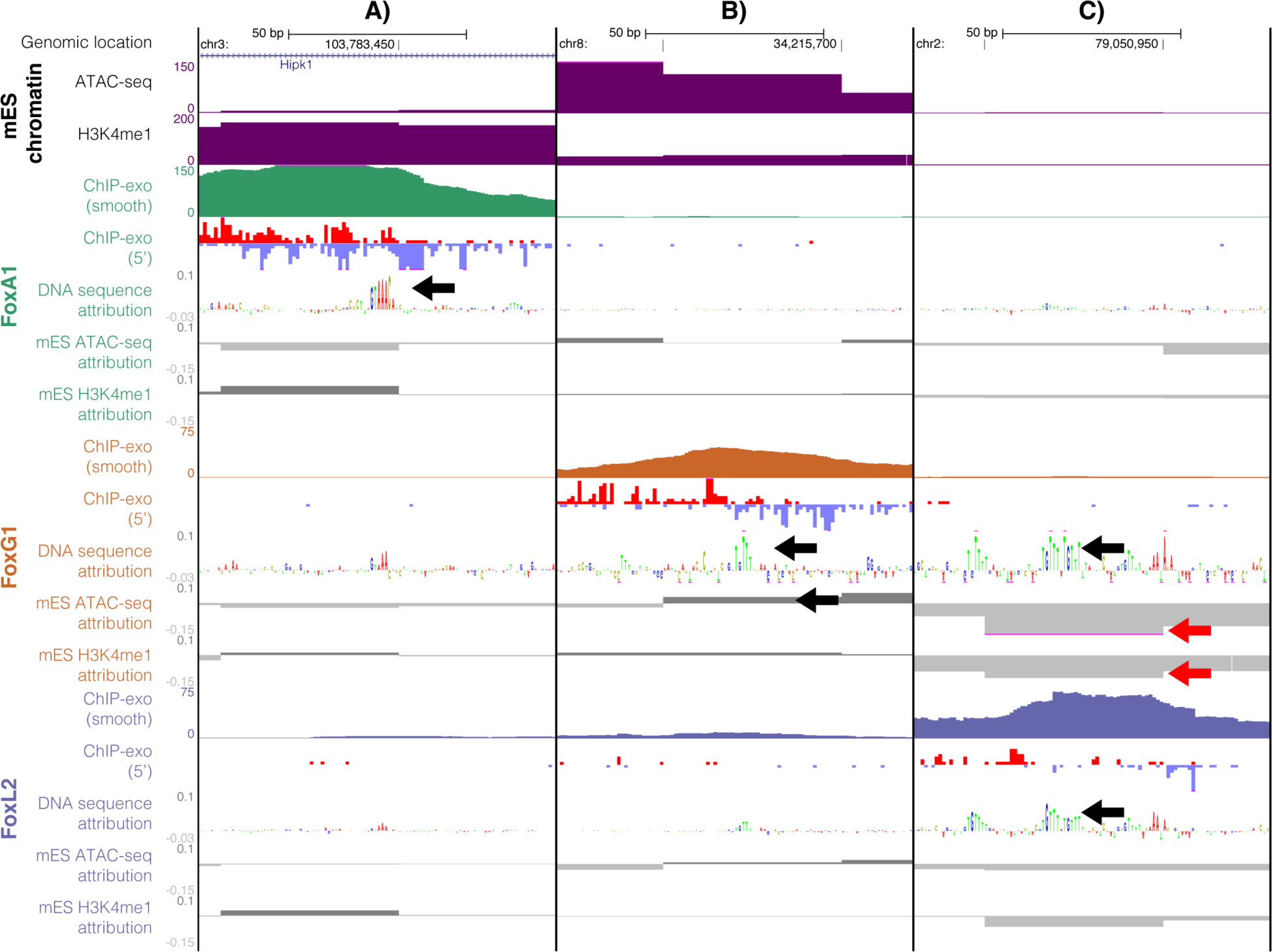
Sequence+chromatin CNN feature attributions explain differential binding at selected Fox TF binding sites. The three UCSC genome browser columns are centered on: **A)** a FoxA1-preferred site at chr3:103783444; **B)** a FoxG1-preferred site at chr8:34215670; and **C)** a FoxL2-preferred site at chr2:79050928 (mm10 coordinates). Each panel shows mES ATAC-seq and mES H3K4me1 ChIP-seq read coverage (50bp resolution). For each of FoxA1, FoxG1, and FoxL2, the browser plots show ChIP-exo read coverage (both an unstranded smoothened read coverage and the stranded ChIP-exo read 5’ positions), DNA sequence attributions, and mES ATAC-seq and H3K4me1 chromatin feature attributions (positive attributions = dark grey above line; negative attributions = light grey below line). In each panel, positive and negative attribution scores referred to in the text are highlighted with black or red arrows, respectively.

### Differential Fox binding produces distinct impacts on chromatin and gene expression

Having demonstrated that Fox paralogs vary in their abilities to bind to relatively inaccessible chromatin, we next asked whether the various Fox paralogs cause chromatin to become more accessible once bound to their target sites. We used ATAC-seq as an indicator of chromatin accessibility, performing experiments in the preexisting mES cells (0hrs), 24hrs after expression of each of the five Fox paralogs, or 24hrs of differentiation with a null control mES cell line. We then plotted ATAC-seq signals over the top 2,000 binding sites of each Fox TF. Sites that overlap a preexisting “active” mES chromatin state are plotted separately from the other sites, which we refer to here as mES “inactive” (**Fig. 7A**, **Fig. 7B**; see Methods for definition of “active” mES states).

**Figure 7:**
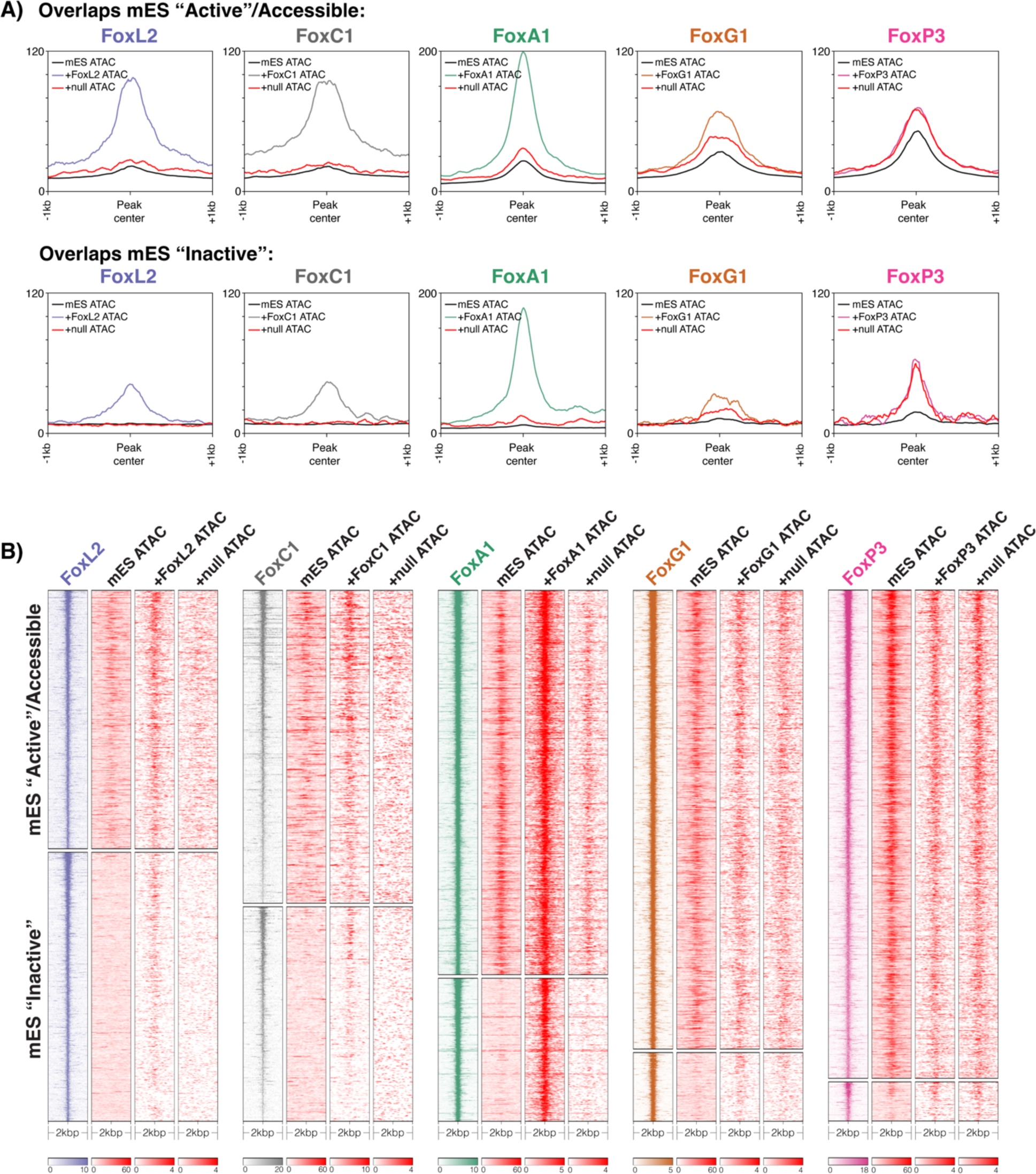
Fox TFs display distinct abilities to open chromatin at their binding sites in mES cells. **A)** Profile plots and **B)** heatmaps for ATAC-seq experiments at the top 2,000 binding sites for each Fox TF. ATAC-seq experiments were performed in mES cells (black line in profile plots), after 24hrs of expression of each Fox TF (colored lines in profile plots), and after 24hrs of differentiation in a null control line (red line in profile plots). Each Fox TF’s binding sites are split into two groups according to their overlap with “active” or accessible regions in the preexisting mES cells (see Methods).

The known pioneer factor FoxA1 causes a substantial increase in ATAC-seq signal at its binding sites, regardless of whether they were “active” or “inactive” in the preexisting mES chromatin environment. FoxL2 and FoxC1 cause a similar increase in accessibility at their binding sites, albeit with a relatively lower gain in accessibility at “inactive” sites when compared with FoxA1. In contrast, FoxP3 and FoxG1 display little if any ability to cause an increase in accessibility at their binding sites. While ATAC-seq signal does increase at FoxP3 and FoxG1 binding sites relative to mES levels, similar gains in accessibility are observed in the null control line at the same sites (**Fig. 7A**, red lines). This suggests that the increase in accessibility at FoxP3 and FoxG1 binding sites occurs due to other TFs that are activated during early differentiation and is not caused by FoxP3 or FoxG1 binding. Thus, the Fox paralogs not only differ in their abilities to bind to relatively inaccessible chromatin, but also in their abilities to open those regions during differentiation.

Next, we asked whether the differential binding activities observed across Fox paralogs have a consequence for downstream gene expression patterns. RNA-seq experiments show that expression of each Fox TF for 24hrs results in unique differential expression patterns (**Fig. 8A**). FoxP3 and FoxG1, the two paralogs that bind to relatively accessible chromatin, produce the most similar transcriptional outputs. The differential expression patterns observed downstream of induced Fox TFs are consistent with previous microarray-based gene expression profiles in the same cell lines [38–40]. Developmentally relevant tissue types were detected in gene set enrichment analyses that focused on the up-regulated TFs downstream of each Fox TF (**Fig. S13**). Amongst other enriched terms, we find endodermal (lung & pancreatic) tissue terms enriched for upregulated TFs downstream of FoxA1, kidney and cornea tissue terms downstream of FoxC1, ovary terms downstream of FoxL2, and neuronal tissue and T-cell terms downstream of FoxG1 and FoxP3. While ectopic Fox TF expression in mES cells is not expected to faithfully recapitulate specific developmental systems, the listed tissue type associations are consistent with known developmental roles of the relevant Fox TFs [28]. Finally, the differential patterns of Fox TF-DNA binding can be linked to the observed differential expression patterns (**Fig. 8B**, **Fig. S14**). For example, the binding site clusters that are preferentially bound by FoxA1 (clusters “A1 (1)” and “A1 (2)” in **Fig. 2B**) are enriched near genes that are specifically up-regulated downstream of FoxA1 (clusters 1-4 in **Fig. 8A**), while FoxG1 has a greater overall association with down-regulated genes, consistent with its hypothesized role as a transcriptional repressor [46] (**Fig. S14**).

**Figure 8:**
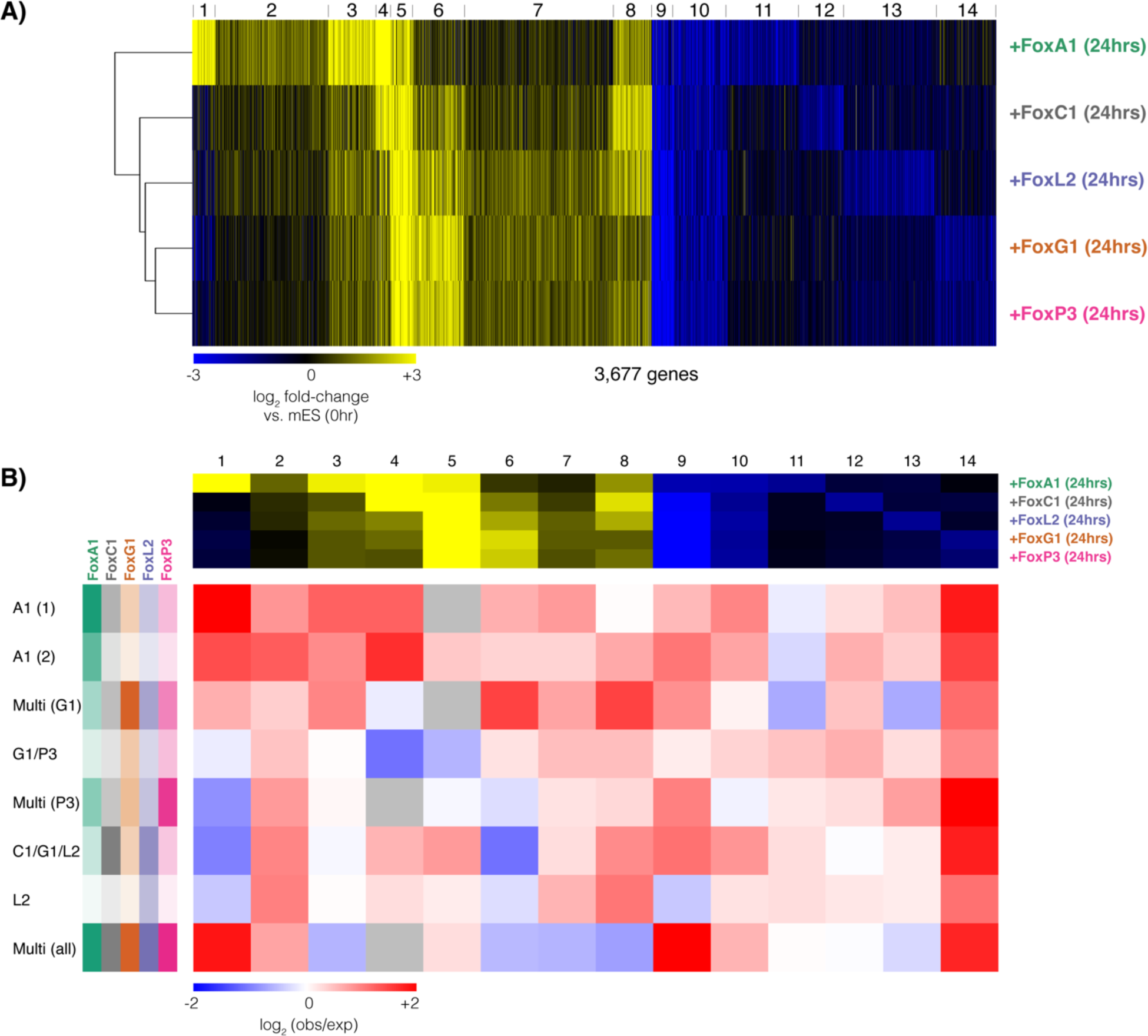
Fox TFs induce distinct transcriptional responses when expressed in mES cells. **A)** K-means clustering of differentially expressed genes detected after 24hrs of Fox TF induction (DESeq2; >2 fold-change, FDR<0.01). **B)** Enrichment/depletion of Fox TF binding sites (clusters defined in Figure 2) located near differentially expressed genes clusters (defined in panel A). Log2 observed/expected ratios are displayed using a blue (depleted) to red (enriched) color shading. At the top of the panel, the average log2 foldchanges (vs. mES 0hr) of genes in each gene expression cluster are displayed for reference. To the left of the panel, the average ChIP-exo read counts of binding sites in each cluster are displayed for reference.

## DISCUSSION

We have demonstrated that paralogous Forkhead transcription factors vary broadly in their genomic binding patterns, even when expressed in the same chromatin environment. Differential Fox TF binding patterns can be partly explained by differences in their intrinsic DNA-binding (sequence) preferences; FoxA1 binds a full TGTTTAC motif whereas the other tested Fox TFs appear to prefer shorter TGTTT motif variants. Our motif analyses also hint that preferences for short tandem repeats of T_n_G sequences may also play a role in specifying binding sites (e.g., as in refs. [47,48]; see **Fig. 2B**, **Fig. S4B**, **Fig. S11**).

Beyond sequence features, our analyses of differential binding patterns using joint sequence & chromatin neural networks and feature attribution techniques suggests that the Fox paralogs also differ in their relative affinities for preexisting chromatin states. Some Fox TFs (e.g., FoxC1 and FoxL2) can bind to sites that lacked chromatin accessibility or other regulatory features in mES cells. Others (e.g., FoxG1 and FoxP3) prefer to bind to regions that already display chromatin accessibility and/or the binding of other TFs in mES cells. The classical pioneer factor FoxA1 mixes both of these binding modes, potentially explaining why its binding sites vary across different cell types despite having documented pioneering activities.

Our work thus strongly implies that Fox TFs should not be assumed to have pioneering activities based solely on structural similarities of the Forkhead domain across the family. Several other Fox paralogs have been proposed to act as pioneer factors during cellular differentiation. For instance, both FoxE1 and FoxO1 have been documented to bind to individual nucleosomal sites [30,31]. FoxF1 was proposed to be a pioneer factor in gastrointestinal stromal tumors based on a loss of ATAC-seq signal at its binding sites in the context of a FoxF1 knock-down [49]. FoxH1 is proposed to bookmark developmental genes during the Maternal-Zygotic Transition [50], while FoxD3 and FoxK2 are proposed to act as early-developmental placeholders for future regulatory events [21,51]. But beyond studies of individual factors, little is known about how pioneering activities vary across Fox TFs. We sought to address this shortcoming by comparing the binding behaviors of the selected Fox paralogs in a common chromatin environment.

While we have shown that differential sequence and chromatin binding preferences guide Fox paralogs to different binding sites, we do not expect that differential binding will explain all divergence in Fox TF regulatory activities. For example, FoxD3 switches from an activator to a repressor during zebrafish neural crest development, suggesting that cell-specific co-factor associations can drive Fox TF regulatory activities [52]. Additionally, a lack of pioneering activities does not necessarily mean that a Fox paralog lacks the ability to serve as a potent developmental regulator. FoxP3 is highly associated with preexisting mES chromatin features in our study and lacks pioneering activity in T-cell development [32]. However, FoxP3 remodels chromatin architecture to rewire enhancer-promoter interactions [53,54], thereby playing a lineage-determining role in T-cell development. Indeed, a recent report showed that FoxP3 can multimerize to form a bridge between two T_n_G repeat sequences [48].

Our observations demonstrate that Fox TFs with similar DNA-binding preferences can have distinct genomic occupancy patterns, depending on their interactions with the preexisting chromatin landscape. We have previously documented a similar phenomenon for a selection of paralogous Hox homeobox transcription factors [55]. In that previous work, Hox TFs with similar DNA-binding preferences were observed to bind to distinct sites, and drive distinct cell fates, when over-expressed in mES-derived progenitor motor neurons. A clear determinant of differential Hox DNA-binding activities was differential associations with sites that were relatively inaccessible in the preexisting chromatin environment. Diverging chromatin affinities may thus be a common strategy by which evolution diversifies the regulatory activities of paralogous TFs that bind to similar motifs.

Efforts to systematically characterize *in vitro* protein-DNA interactions have been hugely valuable in cataloging the intrinsic DNA-binding preferences of TFs. However, given widespread divergence in TF-chromatin association preferences, *in vitro* experiments will not be sufficient to explain regulatory divergence amongst related TFs. We propose that larger-scale efforts to characterize the binding of exogenously expressed TFs in well-characterized chromatin environments will be necessary to fully characterize the relative chromatin affinities of related TFs. Neural network models and feature attribution techniques can then be more broadly applied to understand how interactions between proteins, DNA, and chromatin together determine TF genomic occupancy.

## METHODS

### Fox DNA-binding domain phylogeny

The phylogenetic tree presented in **Fig. S1A** was created by first downloading DNA-binding domain sequences for all Fox paralogs in mouse from cis-bp [34] (version 2.0). We excluded FoxR1 from analysis due to an incomplete DBD sequence in cis-bp. MEGA [56] (version 11.0.9) was used for phylogenetic analysis. Within MEGA, sequences were aligned using Muscle (default settings) and a maximum likelihood tree was constructed with LG substitution model, Gamma distributed rates among sites (5 discrete Gamma categories) and partial deletion (90% cutoff) of gaps and missing data. DNA-contacting residues in **Fig. S1B** are annotated according to ref. [57].

### mES cell lines and culture

Generation of the mES transgenic cell lines used in this study is described in ref. [38]. These cell lines were part of the NIA (National Institute on Aging/NIH) mouse ES cell collection [38–40]. Cell lines are derived from MC1 mouse ES cells (129S6/SvEvTac). Doxycycline-repressible expression constructs containing the relevant TF ORF, C-terminally tagged with His6-FLAG, and a puromycin resistance gene were inserted into the expression-competent Rosa26 locus [38]. Induction of TFs were carried out as described in the original manuscript [38]. In brief, ES cells were cultured in a gelatin-coated dish with DMEM medium containing 15% FBS, 1000 U/ml LIF, 1 mM sodium pyruvate, 0.1 mM MEM non-essential amino acids, 2 mM glutamate, 0.1 mM ß-mercaptoethanol, 0.2µg/ml Doxycycline and 1.0μg/ml puromycin. To induce transgene TFs, Dox was removed by washing three times with PBS at 3 hr intervals and cells were grown in DMEM differentiation medium containing 10% FBS. Chromatin was crosslinked using 1% formaldehyde for 10 minutes.

### ChIP-exo

ChIP-exo was performed following the ChIP-exo 5.0 protocol [58]. Lysis was performed in Farnham cell lysis buffer for 20mins at 4°C. At the 10-min mark, cells were pushed through a 25-gauge needle five times to enhance cellular lysis. Nuclei were then isolated by pelleting at 10,000 rpm for 3 min. Nuclei were resuspended in RIPA buffer (25 million cells to 1 mL of buffer) for an additional 20 min at 4°C and then pelleted again at 10,000 rpm for 3 min. Disrupted nuclei were then finally resuspended in 1× PBS (25 million cells to 1 mL of buffer) and sonicated for 10 cycles (30-on/30-off) in a Diagenode pico. Solubilized chromatin was pre-cleared by centrifugation at maximum speed for 15 min at 4°C before incubating with 10 μL pAG Dynabeads, preloaded with 3 μg of antibody overnight, then sequentially processed through A-tailing, first adapter ligation, Phi29 fill-in, lambda exonuclease digestion, cross-link reversal, second adapter ligation, and PCR followed by size selection for final high-throughput sequencing. Equal proportions of ChIP samples were barcoded, pooled, and sequenced. Illumina paired-end read (40bp read1 and 36bp read2) sequencing was performed on an Illumina NextSeq 500. The 5′ end of read1 corresponds to the exonuclease stop site, located ∼6 bp upstream of a protein-DNA cross-link. read2 serves two indirect functions: to provide added specificity to genome-wide mapping and to remove PCR duplicates. The following antibodies were used: anti-FLAG-M2 (F1804 Sigma, lot SLBW5142), anti-IgG (i5381 Sigma, lot SLBQ7097V), anti-Foxa1 (Abcam ab23738, lot GR3241520-1), anti-Ascl1 (ab74065 Abcam, lot GR3209594-1). Null control ChIP-exo experiments were performed using the anti-FLAG-M2 antibody in MC1 mouse ES cells that did not carry a Dox-inducible expression construct.

### Choosing ChIP-exo time-point and antibody

To assess an appropriate time-point for capturing induced TF binding activities, we performed an initial time course of ChIP-exo experiments for FoxA1 after initiating transgene expression. The numbers of binding sites resulting from ChIP-exo does not become robust until 24 hrs of expression (**Table S1**) and we did not see substantial differences in FoxA1 binding between 24hrs and 48hrs (**Fig. S2**). We confirmed these trends by performing a similar ChIP-exo timecourse on another known pioneer TF, Ascl1 (**Table S1**). We also confirmed that the binding sites captured by targeting the FLAG epitope tag on the FoxA1 transgene are not substantially different from those captured via a native FoxA1 antibody (**Fig. S2**). Subsequent experiments thus focused on the 24hr timepoint (as it represents the earliest timepoint where we can capture robust Fox TF binding activities in this system) and the FLAG antibody (since we can target it consistently across all selected Fox TFs). To minimize batch effects, additional replicates of FoxA1 (anti-FLAG) ChIP-exo experiments were performed at the 24hr timepoint alongside the ChIP-exo experiments for the other Fox TFs. Analysis of the new experiments accounts for the discrepancy in FoxA1 peak numbers between Table S1 and Fig. 1.

### ChIP-exo data processing

BWA mem (version 0.7.17-r1188) [59] was used to align paired-end ChIP-exo reads against the mm10 reference genome using the following arguments: “-t 8 -v 1 - T 30 -h 5”. Picard MarkDuplicates (version 2.7.1) was used to mark duplicate read pair alignments and then samtools view with arguments “-h -b -f 0×1 -F 0×404” was used to filter out duplicated read pairs. While read pairs were used for alignment and deduplication purposes, only read1 alignment information from the deduplicated alignments was retained for downstream analysis (read1 corresponds to the ends of the immunoprecipitated DNA fragments that are subject to exonuclease digestion, whereas read2 information has the same resolution as ChIP-seq).

Initial peak-finding was performed for each ChIP-exo experiment with ChExMix (version 0.52) with the following settings: “--threads 8 --scalewin 1000 --mememinw 6 --mememaxw 18 -- q 0.05 --epsilonscale 0.5”, excluding ENCODE blacklist regions [60], and using MEME (version 4.11.3) [61] internally to discover motifs associated with binding subtypes. To encourage consistency in shared peak coordinates across Fox TFs, we re-ran ChExMix in a mode where the possible binding subtype motifs are pre-initialized and remain fixed during training, using settings: “--threads 8 --scalewin 1000 --q 0.05 --epsilonscale 0.75 --motfile $motifs – noupdateinitmotifs”. The motif set provided as “$motifs” contain the MEME-discovered subtype motifs from the most populous binding subtypes in the initial ChExMix runs. Note that for a given Fox TF, ChExMix models the data from each biological replicate separately but produces a single set of peaks that are consistent across replicates.

Histograms of distances between ChIP-exo peaks and nearest TSSs were created using GREAT (version 4.0.4) [62]. Associations between Fox TF peaks and repeat families were created by overlapping with RepeatMasker annotations using bedtools intersect -u. Only repeat families with at least a 1% overlap were used in the related figure. Expected overlap rates were calculated by randomly sampling the same number of genomic locations as elements in a given repeat family (ignoring ENCODE blacklist regions [60]), repeated 100 times. Log_2_ observed/expected rates were plotted in R using ggplot2.

### Fox binding site differential binding and clustering

For the purposes of assessing differential binding and clustering Fox TF binding patters, we first combined the top 5,000 peaks from each Fox TF (ranked by ChIP-exo enrichment level), merging sites which were within 20bp of each other. To calculate replicate-wise and condition-wise ChIP-exo signal enrichment for each Fox TF across the merged set of sites, we used custom Java code (org.seqcode.projects.sequtils.EnrichmentTester2 in https://github.com/seqcode/seqcode-dev) with the following settings: “--q 0.01 --minfold 2 --win 200 --simple --fixedpb 10000 --joinwin 1 -- medianscale”. EdgeR (R version 4.1.0, edgeR version 3.34.0) [63] was used internally in the code to normalize between conditions and to determine differentially bound sites across conditions (fold>1.5, q<0.05). Normalized signal values (calculated using edgeR scaling factors) were K-means clustered in R, where K was varied from 7 to 20. The final value K=10 was chosen by subjectively trying to minimize number of small clusters while preserving interesting differences between clusters. K=10 provided greater distinction between C1-preferred, G1-preferred, P3-preferred and the other clusters. K=11 started showing subdivision of existing cluster classes without additional informative clusters. Within the K=10 cluster set, two pairs of clusters were merged due to similar enrichment patterns across TFs, thus yielding 8 final clusters.

DeepTools (version 3.5.1) [64] was used to plot ChIP-exo signal intensity across the clustered binding sites. Bigwig files were created using the DeepTools bamcoverage utility with binsize 1 and scaling factors calculated from edgeR above. The DeepTools computeMatrix utility was run with the following options “reference-point --referencePoint center -a 1000 -b 1000 -p 8 --missingDataAsZero --sortRegions keep”

### Motif analysis

MEME-ChIP (version 5.3.3) [61] was run on 100 base pair windows around the sites in each cluster with settings “-meme-nmotifs 5 -meme-mod zoops -minw 6 -maxw 20 -db JASPAR/JASPAR2018_CORE_vertebrates_non-redundant.meme”. SeqUnwinder (version 0.1.5) [42] was run in multi-class mode, where each of the K-means clustered binding sites was labeled according to cluster membership. SeqUnwinder settings were: “--threads 10 -- makerandregs --win 150 --mink 4 --maxk 5 --r 10 --x 3 --a 400 --hillsthresh 0.1 -- memesearchwin 16”, where MEME (version 5.3.3) was used internally to build motifs from discriminative k-mer features.

### Comparison with mES TF binding and chromatin signals

We compared the Fox TF binding sites with several previously published mES histone modification ChIP-seq, TF ChIP-seq, and ATAC-seq experiments (database identifiers listed in **Table S2**) [44,65–68]. Locations of enriched domains in the mES histone modification and ATAC-seq experiments were determined using a custom domain finder algorithm (org.seqcode.projects.seed.DomainFinder in https://github.com/seqcode/seqcode-core) using the following arguments: “--threads 4 --binwidth 50 --binstep 25 --mergewin 200 --poisslogpthres -5 --binpthres 0.05. This has the effect of finding domains that are significantly enriched over control experiment levels with p<0.05. The top 2,000 binding sites from each Fox TF (ranked by ChIP-exo enrichment level) were compared against each histone mark’s domains using bedtools window -w 200 (version 2.30).

ChromHMM was run using default settings where we first binarized bed files using the BinarizeBed command and then learned a chromatin state model using the learnModel subcommand (choosing 15 states). The top 2,000 binding sites from each Fox TF (ranked by ChIP-exo enrichment level) were again compared against each chromatin state’s locations using bedtools window -w 200 (version 2.30).

Regions of “active” mES chromatin (e.g., for **Fig. 7**) were defined as regions that were covered by ChromHMM states “EnhG”, “EnhLo”, “Enh”, “EnhPois”, “Tss”, “TssBiv” (labeled according to the state names in Fig. 3B), or regions covered by an mES ATAC-seq enriched domain.

### ATAC-seq

ATAC-seq experiments followed the Omni-ATAC-seq protocol [69]. For each experiment, 50,000 cells were harvested. Cells were dissociated in 50 μl ATAC-Resuspension Buffer (10 mM Tris (pH 7.4), 10 mM NaCl, 3 mM MgCl2) with 0.1% NP-40, 0.1% Tween-20, and 0.01% Digitonin, incubated on ice for 3 mins, and then suspended in 1 mL ATAC-Resuspension Buffer with 0.1% Tween-20 and centrifuged for 10 min at 4°C and 500g. The pellet was resuspended in transposition mixture (25 μl 2x TD buffer, 2.5 μl transposome (Tn5 + oligo DNA), 16.5 μl PBS, 0.5 μl 1% digitonin, 0.5 μl 10% Tween-20, 5 μl H_2_O), followed by incubation at 37°C for 30mins. The reaction was cleaned up with AMPure XP beads. Transposed fragments were PCR amplified for a number of PCR cycles determined from one-third of the maximum fluorescence measured by quantitative PCR reaction with 1 × SYBR Green (Invitrogen), custom-designed primers [70], and 2 × NEB MasterMix (New England Labs, M0541). The library was cleaned up with AMPure XP beads and the fragment length distribution of the library was determined using a Bioanalyzer High Sensitivity chip. The libraries were sequenced on an Illumina NextSeq 2000. ATAC-seq experiments were performed in duplicate. ATAC-seq heatmaps and profile plots were created using DeepTools (version 3.5.1) [64].

### RNA-seq

RNA-seq experiments started with 1 μg of total RNA. The NEBNext Poly(A) mRNA Magnetic Isolation Module (NEB #E7490) was used to enrich for polyA mRNA. Strand-specific RNA-seq libraries were then prepared using the NEBNext® Ultra II Directional RNA Library Prep Kit. RNA-seq experiments were performed in duplicate. Illumina sequencing was performed on an Illumina NextSeq 500. One set of replicates was sequenced paired-end (40bp read1 and 36bp read2), while the other was sequenced single-end (75bp).

### RNA-seq analysis

Gene expression was quantified by mapping the RNA-seq fastq files to the mouse mm10 GENCODE transcriptome (version 23) [71] using Salmon (version 1.4.0) [72] with the arguments ‘-l A --gcBias --validateMappings’. Differential gene expression analysis was performed using DESeq2 (version 1.30.1) [73]. To account for the fact that there were single-end and paired-end replicates for the induced FoxC1, FoxG1, FoxL2, and FoxP3 cell-line at the 24hr time point, we used a design formula ‘readtype + condition’. For the 0hr time-point, we treated all cell-lines as the same condition, whereas for the 24hr time-point, the condition was the concatenated cell-line and time-point. To plot the heatmap of RNA-seq expression, we first used DESeq2 to calculate the log_2_ foldchange of expression between the 24hr time point for each cell-line and the 0hr condition. We selected genes with an absolute foldchange of >2 and an FDR<0.01 in at least one condition, resulting in 3,710 genes that were taken forward for clustering. The log_2_ foldchange values across all 5 Fox TFs were clustered in R (version 3.6) using the kmeans function with k=16. Two clusters containing fewer than 30 genes were removed from consideration, leaving 3,677 clustered genes. Associations between sets of binding sites and sets of genes were performed using GREAT command line tools (version 4.0.4) [62]. Specifically, we defined regulatory domains for all GENCODE genes using GREAT tools createRegulatoryDomains with arguments “basalPlusExtension -maxExtension=100000 - basalUpstream=5000 -basalDownstream=1000”, overlapped the peak locations within set of binding sites with gene regulatory domains from each set of genes using bedtools intersect -u, performed the same overlaps with randomly selected genomic locations matched in size to each binding site set (10 repetitions), and assessed overlap significance using GREAT tools calculateBinomialP. Gene set enrichment analysis of tissue-specific terms was performed using EnrichR [74], focusing on gene sets provided by Tabula Muris [75] and the Mouse Gene Atlas [76].

### Convolutional neural network training, validation, and test set preparation

The length of each input region is set to 500bp to capture the local sequence and chromatin context. Three chromosomes were held out for validation (chr11) and testing (chr2, chr17). Positive samples for each Fox TF are defined using ChExMix peak summits as determined above. Training set samples randomly shifted the 500bp window around the peak summit (within a range of ±225bp) to avoid the CNN model only focusing on the center region of the input. Each ChExMix peak summit was shifted multiple times to augment the positive training examples. The number of shifting iterations varied for each Fox TF, due to the varying numbers of peaks. Specifically, we shifted training peaks the following numbers of times: FoxA1 = 5; FoxC1 = 125; FoxG1 = 5; FoxL2 = 25; FoxP3 = 25. For validation and test sets, positive samples are 500bp windows centered on the ChExMix summit. Negative samples are defined using randomly sampled 500bp windows, excluding any window that overlaps a Fox TF ChExMix peak or an ENCODE blacklist region [60]. The negative training set size was set to match the positive training set size. After obtaining positive and negative sets for all five Fox TFs, we combined all of them into a “meta” dataset for training, validation, and testing, serving the purpose of showing models more regions that exhibit diverse Fox family factor binding preference. For example, a window that is bound only by FoxA1 would be used as a positive training sample for the FoxA1 training task, and as a negative sample for the other Fox TF training tasks. The model accepts a one-hot-encoded DNA sequence and a chromatin signal array as inputs. Given a 500bp input region, one-hot-encoded DNA was prepared by mapping each nucleotide according to the dictionary: {A=[1,0,0,0], T=[0,1,0,0], G=[0,0,1,0], C=[0,0,0,1]}, resulting in an array of shape (4, 500). A selection of mES chromatin experiments was sourced from previous publications (database identifiers listed in **Table S3**) [77–79]. ChIP-seq and ATAC-seq reads were processed similarly to ChIP-exo above, but using bowtie2 [80] (version 2.3.5) with default settings. Per-base coverage information was extracted from each chromatin track’s bigwig file, transformed into a standardized read count (z-score), and stacked into an array of shape (# *chromatin tracks*, 500). One-hot-encoded DNA sequence and chromatin arrays are integrated to become a single array of shape (4 + # chromatin tracks, 500) that is input into the CNN. The prediction target of the model is the relevant Fox TF’s log transformed ChIP-exo read count overlapping the region.

### Convolutional neural network architecture and training

Our CNNs use a convolutional tower architecture inspired by the VGG architecture [81]. The model consists of three parts:

1. The first part of the model is a single 1-D convolutional layer that includes *n* filters of kernel size 25 and stride 1, which serves the purpose of extracting motif-like sequence features and local chromatin patterns.
2. The middle part is a convolutional tower composed of four 1-D convolutional layers, where each includes filters of kernel size *w* and stride 2. The number of convolutional filters at each layer in convolutional tower is computed as:

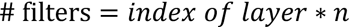
3. A convolutional layer that includes only one filter of kernel size 1 and stride 1 reduces the dimensionality of the tensor. The reduced 1D output is then passed through *k* fully connected layers of *f* output features. Finally, the last dense layer outputs the prediction of TF binding strength.

The full list of operators included in the model is listed in **Table 1**.

**Table 1:**
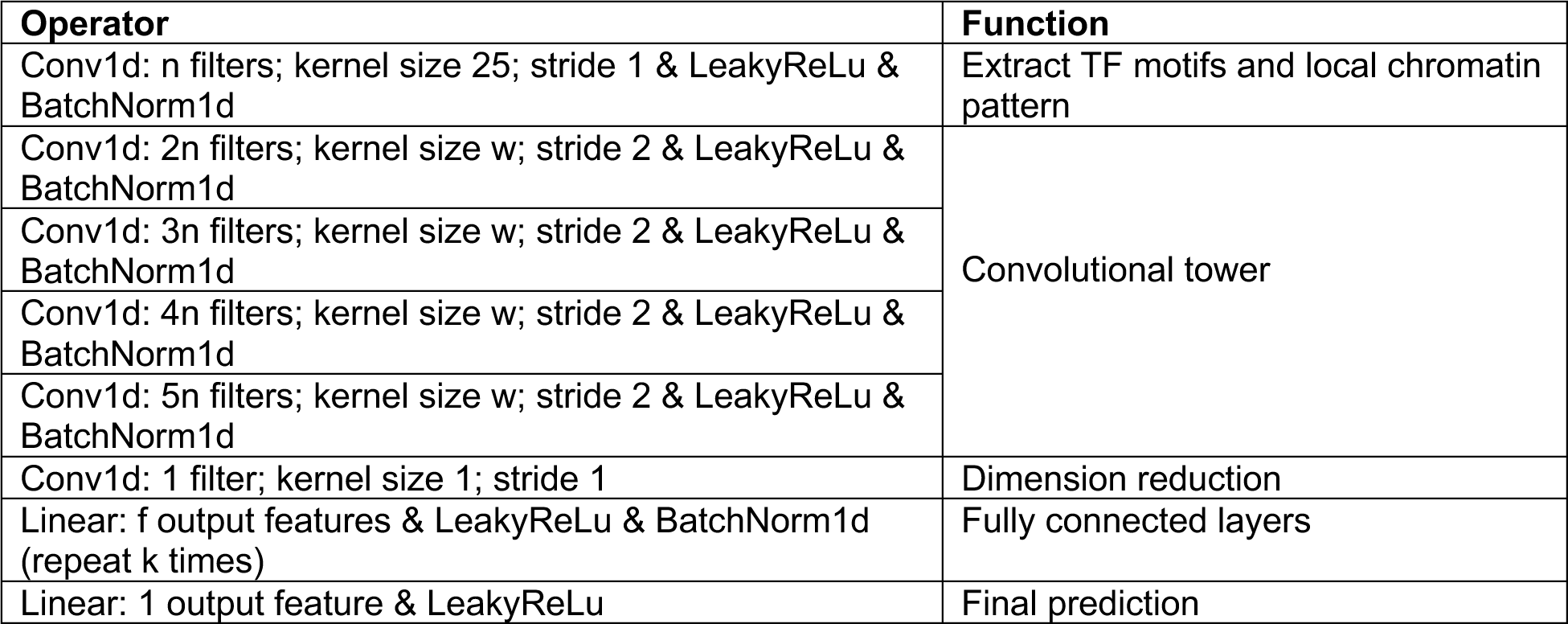
Architecture of the CNN models.

The CNNs were implemented on PyTorch [82] (version 1.11.0) under the PyTorchLightning framework (version 1.7.4). The AdamW optimizer (*learning rate = lr, betas* = [0.9, 0.999], *eps* = 1*e* – 08, *weigt_decay* = 0.01) and Exponential Learning Scheduler (*multiplicative factor* = *gamma*) were used for model training. The hyperparameters *n, w, f, k, lr, gamma* were tuned separately for each Fox TFșfs CNN (sequence+chromatin variant). An Asynchronous HyperBand scheduler implemented by Ray [83] (version 2.0.0) was employed to perform hyperparameter searching for each model. Configurations giving the best performance for each Fox factor are listed in **Table 2**. The same hyperparameters are used for the matching sequence-only CNNs.

**Table 2:** Hyperparameter values yielding the best CNN performance for each Fox TF.

| Factor | # conv. filters n | conv. kernel size w | dense. output features f | dense. repeat k times | batch size | learning rate $lr$ (AdamW) | $gamma$ (Exponential Learning Scheduler) |
| --- | --- | --- | --- | --- | --- | --- | --- |
| FoxA1 | 32 | 5 | 64 | 4 | 256 | 0.0002 | 1.0 |
| FoxC1 | 64 | 3 | 128 | 4 | 256 | 0.00006 | 1.0 |
| FoxG1 | 32 | 3 | 128 | 3 | 256 | 0.00015 | 0.9 |
| FoxL2 | 32 | 5 | 128 | 3 | 256 | 0.00015 | 0.9 |
| FoxP3 | 32 | 5 | 128 | 4 | 256 | 0.00005 | 1.0 |

Model training was performed on two Nvidia P100 GPUs under single precision. Mean Squared Error (MSE) is used as the loss function. The training was early stopped if validation loss was not improved for 10 epochs. Test dataset performance was then evaluated on trained models.

### Convolutional neural network feature attribution

The DeepLiftShap [84,85] implementation from Captum [86] (version 0.5.0) was used to interpret the contribution of the inputs to model predictions. For each input region, di-nucleotide shuffling was performed 10 times on the DNA sequence to get the DNA background set, while a zero array is employed as the background for the standardized chromatin signal array. Modisco-lite [87] (version 2.0.5) was run on the DeepLiftShap hypothetical importance scores against cisBP [34] motifs in human and mouse to extract high-impact global DNA sequence patterns.

## Data and code availability

ChIP-exo, ATAC-seq, and RNA-seq data are available from GEO under accession GSE244411. UCSC genome browser tracks for ChIP-exo data and CNN feature attributions are available from: https://github.com/seqcode/iFox-mES. CNN dataset preparation and training scripts are also available on GitHub: https://github.com/seqcode/iFox-mES.

## Acknowledgements

This work was supported by NIH NIGMS R35-GM144135 (to S.M.) and NIH NIGMS R01-GM125722 (to S.M. and B.F.P). Work by T.A. and M.S.H.K was supported by JSPS KAKENHI Grant Number JP20H05395. The authors thank Dr. Cheryl Keller of Penn State’s Genomic Research Incubator for DNA sequencing support and advice on ATAC-seq. We also thank members of the Center for Eukaryotic Gene Regulation at Penn State for helpful feedback and guidance.

