## Supplemental Figures and Tables for "Joint sequence & chromatin neural networks characterize the differential abilities of Forkhead transcription factors to engage inaccessible chromatin"

### SUPPLEMENTAL FIGURES & LEGENDS

A)

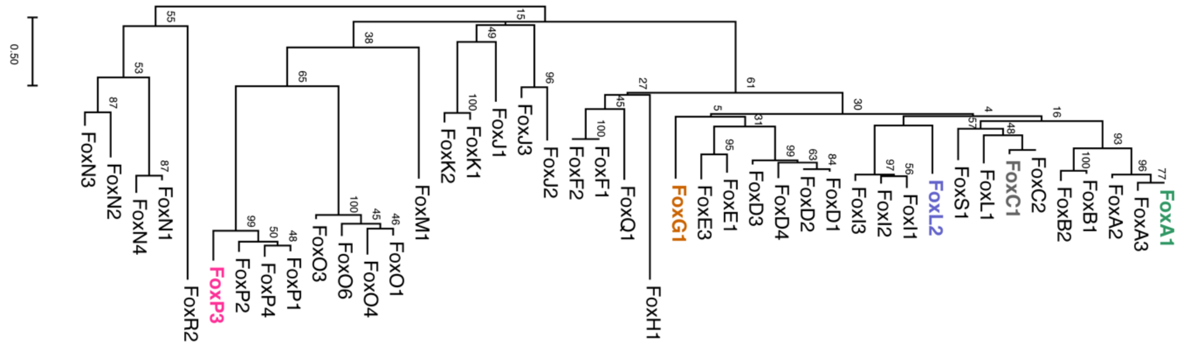

B)

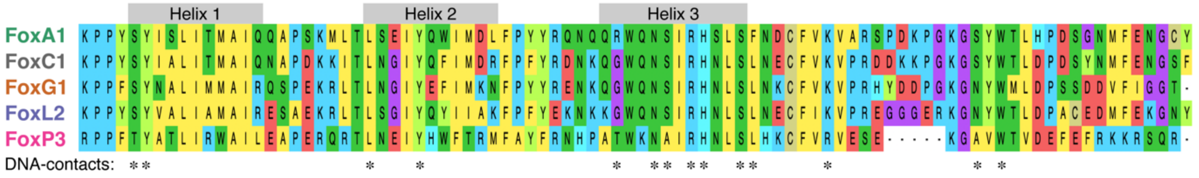

C)

**Identity in aligned DNA-binding domains (%)**

| | FoxA1 | FoxC1 | FoxG1 | FoxL2 | FoxP3 |
|---|---|---|---|---|---|
| FoxA1 | 100% | 73% | 62% | 57% | 39% |
| FoxC1 | 73% | 100% | 69% | 64% | 40% |
| FoxG1 | 62% | 69% | 100% | 65% | 43% |
| FoxL2 | 57% | 64% | 65% | 100% | 38% |
| FoxP3 | 39% | 40% | 43% | 38% | 100% |

D)

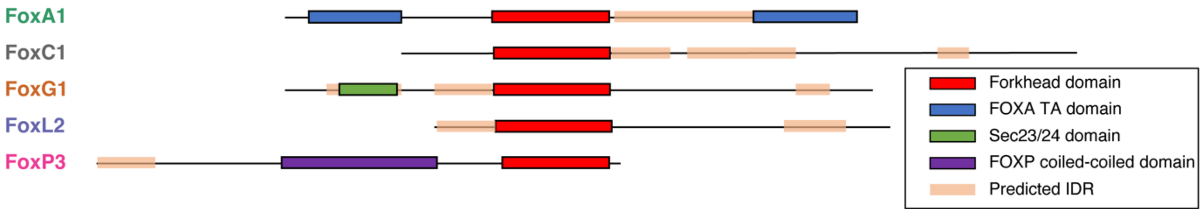

**Figure S1:** Overview of relationships between selected mouse Fox TFs (the five Fox TFs examined in this work are color-coded as shown). **A)** Maximum likelihood tree constructed using aligned Fox TF DNA-binding domains. Aligned sequences are the Forkhead DNA-binding domains for each gene as recorded in the cis-bp database. Bootstrap values showing support for particular branching patterns are displayed at branching points. **B)** Amino acid multiple sequence alignment of the Forkhead DNA-binding domains of the five selected Fox TFs. Structural domains and DNA-contacting residues annotated by ref. [55]. **C)** Amino acid identity rates between Fox TF DNA-binding domains, inferred from the alignment in panel B). **D)** Cartoon representation of the Ensembl-annotated domains in the five selected Fox TFs.

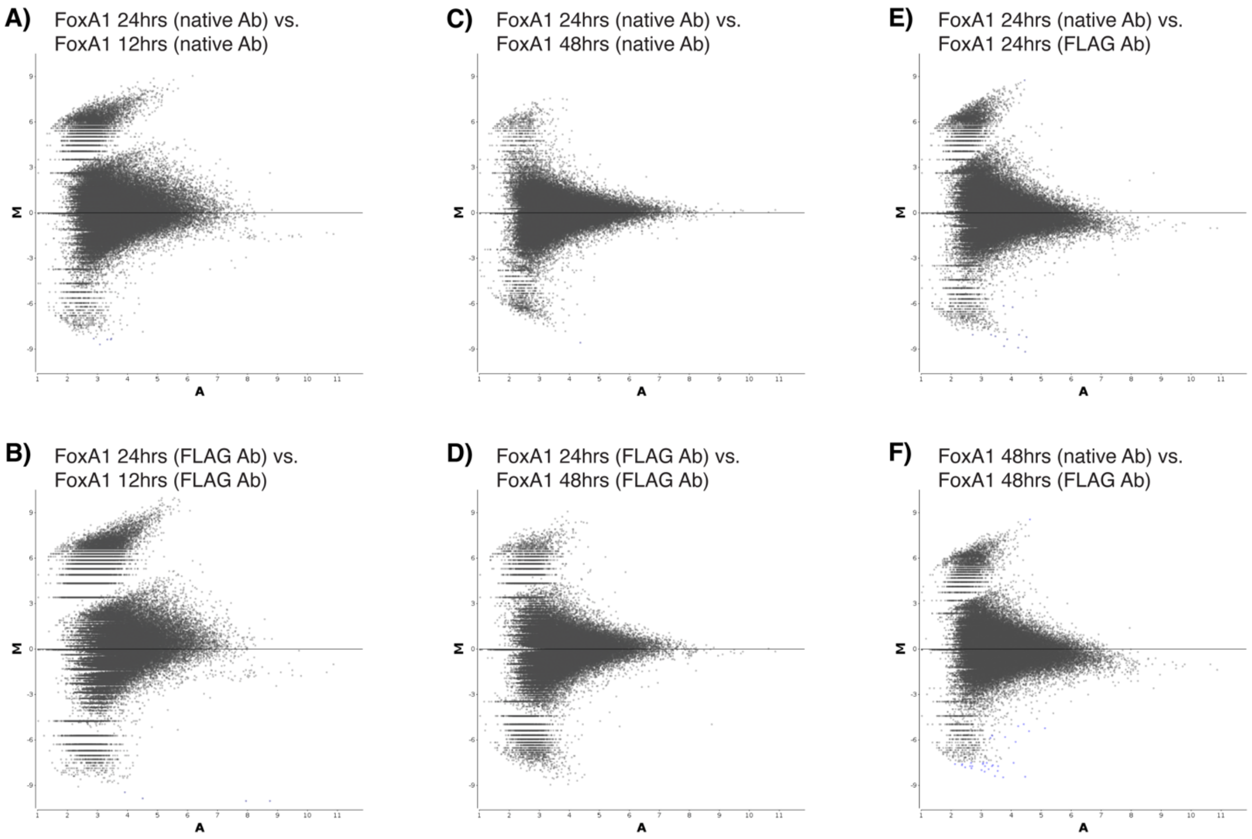

**Figure S2:** MA-plots comparing ChIP-exo read counts across selected pairs of FoxA1 ChIP-exo experiments. The dots in each plot represent the ChIP-exo read counts in 200bp windows around ChExMix-defined peaks merged across the relevant pair of datasets. EdgeR is used to normalize and compare the counts across datasets (see Methods). The y-axes (M) plot the log-foldchange across datasets, and the x-axes (A) plot the average readcount across datasets. Each panel compares a different pair of FoxA1 experiments based on the hours after FoxA1 induction (12hrs, 24hrs, or 48hrs), and/or the antibody used for ChIP-exo (native FoxA1 antibody or anti-FLAG antibody).

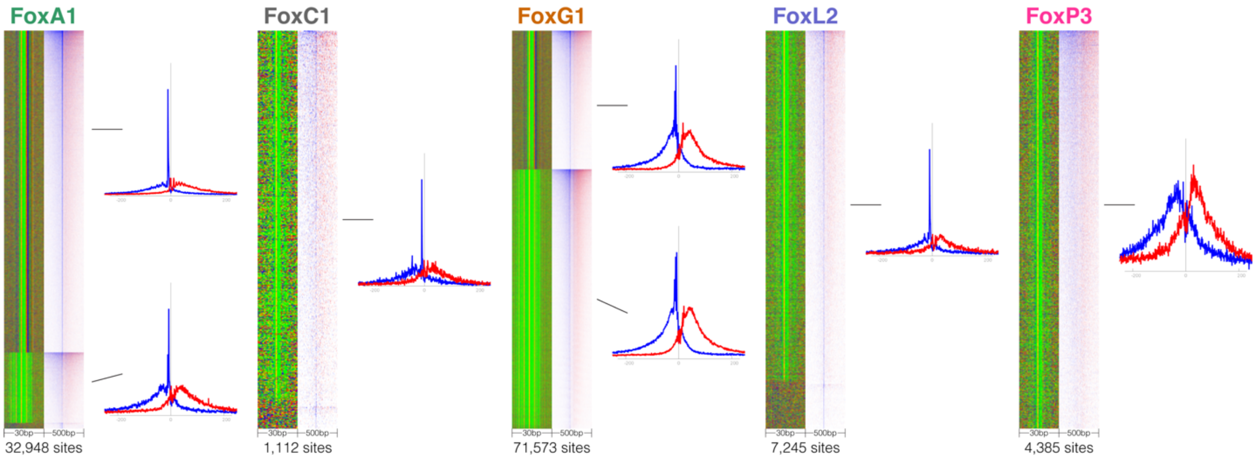

**Figure S3:** ChExMix-defined subtypes for each Fox TF ChIP-exo experiment, as shown in Figure 1, but plotting the profile view of the ChIP-exo read distributions associated with each subtype.

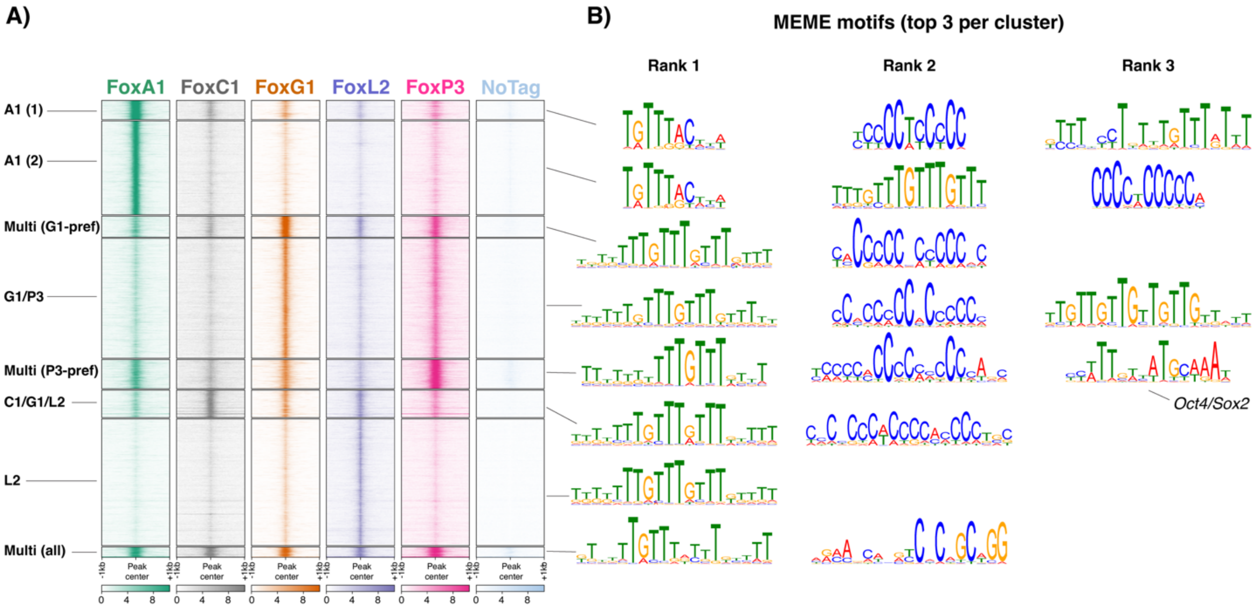

**Figure S4:** Extended version of Figure 2B. **A)** Unstranded ChIP-exo 5' read enrichment plotted at clusters of Fox TF binding sites as determined by K-means clustering. ChIP-exo enrichment from a null control cell line is shown for reference (NoTag). **B)** The top 3 motifs discovered by MEME in each cluster of Fox TF binding sites.

A)

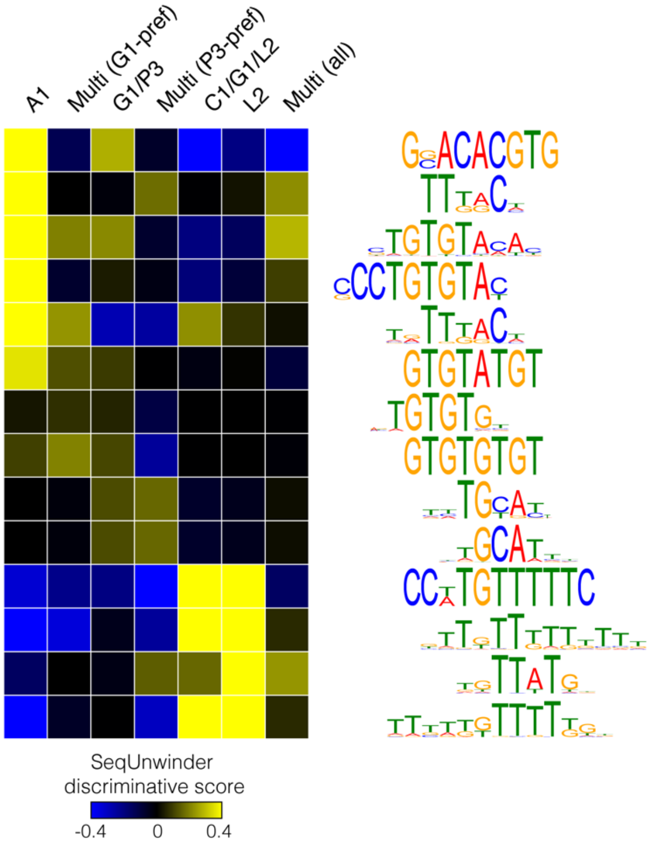

B)

| A1 | Multi (G1-pref) | G1/P3 | Multi (P3-pref) | C1/G1/L2 | L2 | Multi (all) |
|---|---|---|---|---|---|---|
| 5% | 2% | 3% | 3% | 2% | 3% | 4% |
| 40% | 27% | 23% | 25% | 24% | 24% | 23% |
| 53% | 29% | 27% | 27% | 22% | 26% | 24% |
| 41% | 26% | 24% | 26% | 21% | 22% | 29% |
| 70% | 32% | 29% | 34% | 25% | 27% | 34% |
| 17% | 19% | 15% | 8% | 11% | 13% | 11% |
| 45% | 37% | 34% | 28% | 28% | 31% | 33% |
| 22% | 23% | 18% | 11% | 13% | 14% | 16% |
| 25% | 20% | 24% | 27% | 14% | 19% | 18% |
| 21% | 16% | 21% | 29% | 14% | 16% | 15% |
| 18% | 25% | 23% | 21% | 27% | 25% | 30% |
| 35% | 66% | 63% | 44% | 73% | 68% | 55% |
| 16% | 42% | 39% | 23% | 47% | 50% | 30% |
| 27% | 60% | 55% | 35% | 67% | 62% | 47% |

C)

**SeqUnwinder class-wise performance metrics (stratified cross-validation)**

| Class | TP Rate | FP Rate | Precision | Recall | auROC | auPRC |
|---|---|---|---|---|---|---|
| A1 | 0.53 | 0.11 | 0.41 | 0.53 | 0.83 | 0.47 |
| Multi (G1-pref) | 0.26 | 0.10 | 0.28 | 0.26 | 0.69 | 0.24 |
| G1/P3 | 0.15 | 0.08 | 0.21 | 0.15 | 0.63 | 0.18 |
| Multi (P3-pref) | 0.34 | 0.11 | 0.31 | 0.34 | 0.74 | 0.29 |
| C1/G1/L2 | 0.32 | 0.10 | 0.32 | 0.32 | 0.73 | 0.30 |
| L2 | 0.30 | 0.11 | 0.28 | 0.30 | 0.72 | 0.28 |
| Multi (all) | 0.31 | 0.09 | 0.33 | 0.31 | 0.71 | 0.30 |
| Random | 0.54 | 0.06 | 0.54 | 0.54 | 0.90 | 0.58 |

**Figure S5:** SeqUnwinder multi-class discriminative motif-finding analysis on clusters of Fox TF binding sites displayed in Figure 2 (merging the two FoxA1-preferred clusters). **A)** Discovered motifs and the SeqUnwinder scores associating each motif with the relevant clusters of binding sites. **B)** Percentages of sites in each cluster that contain (by FIMO motif scanning) each of the motifs displayed in panel A). **C)** SeqUnwinder multi-class performance metrics in each cluster of sites, including true positive (TP) rates, false positive (FP) rates, precision, recall, area under receiver-operator curve (auROC), and area under precision-recall curve (auPRC).

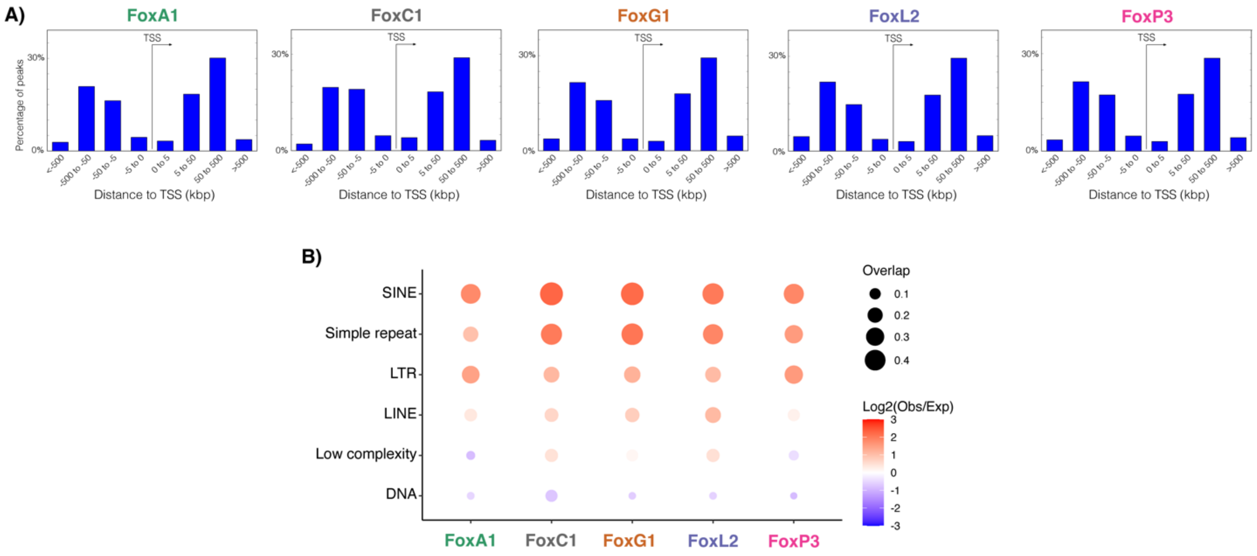

**Figure S6:** Relationships between Fox TF binding sites and genome annotations. **A)** Histograms of distances between Fox TF binding sites and annotated TSSs. **B)** Relationships between Fox TF binding sites and repeat classes, where size of circle reflects the fraction of sites overlapping with the repeat class and color shading reflects the log observed/expected enrichment of overlap.

**A)**

Pre-existing mES TFBS

| | FoxL2 | FoxC1 | FoxA1 | FoxG1 | FoxP3 |
|---|---|---|---|---|---|
| CTCF | 1.4 | 1.1 | 1.1 | 1.1 | 1.4 |
| cMyc | 1.3 | 4.1 | 3.3 | 0.0 | 0.0 |
| Zfx | 1.1 | 2.6 | 3.2 | 2.2 | 3.3 |
| nMyc | 2.0 | 2.3 | 2.4 | 2.4 | 3.2 |
| E2f1 | 1.9 | 3.5 | 4.0 | 4.0 | 4.7 |
| p300 | 2.9 | 3.4 | 5.3 | 5.1 | 6.1 |
| Esrrb | 3.2 | 3.9 | 5.3 | 4.8 | 5.8 |
| Tcfcp2l1 | 3.2 | 3.5 | 5.4 | 4.9 | 5.9 |
| Znf384 | 3.7 | 4.1 | 4.8 | 4.8 | 5.6 |
| Klf4 | 2.7 | 3.7 | 6.3 | 5.4 | 6.4 |
| MafK | 4.0 | 4.1 | 6.0 | 5.9 | 6.9 |
| Stat3 | 5.2 | 0.0 | 7.4 | 6.8 | 8.4 |
| Nr5a2 | 5.6 | 5.3 | 8.0 | 7.9 | 9.3 |
| Oct4 | 5.5 | 5.4 | 8.1 | 8.1 | 9.4 |
| Sox2 | 6.0 | 6.1 | 8.6 | 8.5 | 9.6 |
| Nanog | 6.1 | 6.3 | 8.6 | 8.4 | 9.6 |
| FoxD3 | 7.2 | 7.7 | 9.3 | 9.1 | 9.7 |

log2(overlap enrichment over random)

**B)**

| | FoxL2 | FoxC1 | FoxA1 | FoxG1 | FoxP3 | Random |
|---|---|---|---|---|---|---|
| CTCF | 0.4% | 0.4% | 0.3% | 0.3% | 0.4% | 0.16% |
| cMyc | 0.0% | 0.4% | 0.2% | 0.1% | 0.0% | 0.02% |
| Zfx | 0.2% | 0.7% | 1.1% | 0.5% | 1.2% | 0.12% |
| nMyc | 0.1% | 0.2% | 0.2% | 0.2% | 0.3% | 0.04% |
| E2f1 | 1.2% | 3.7% | 5.3% | 5.1% | 8.3% | 0.32% |
| p300 | 4.0% | 5.7% | 21.4% | 17.8% | 35.4% | 0.53% |
| Esrrb | 4.1% | 7.1% | 18.0% | 12.8% | 25.7% | 0.47% |
| Tcfcp2l1 | 1.7% | 2.2% | 8.0% | 5.6% | 11.2% | 0.19% |
| Znf384 | 3.4% | 4.3% | 7.1% | 7.1% | 13.0% | 0.26% |
| Klf4 | 0.3% | 0.7% | 4.3% | 2.3% | 4.6% | 0.05% |
| MafK | 2.0% | 2.2% | 8.3% | 7.9% | 15.9% | 0.13% |
| Stat3 | 0.2% | 0.1% | 1.1% | 0.8% | 2.3% | 0.01% |
| Nr5a2 | 0.4% | 0.4% | 2.2% | 2.1% | 5.6% | 0.01% |
| Oct4 | 0.7% | 0.6% | 4.2% | 4.0% | 9.8% | 0.01% |
| Sox2 | 1.7% | 1.8% | 9.7% | 9.1% | 19.7% | 0.03% |
| Nanog | 1.4% | 1.6% | 7.8% | 6.7% | 15.5% | 0.02% |
| FoxD3 | 7.4% | 10.0% | 29.9% | 27.2% | 40.9% | 0.05% |

% overlap between peaks

**Figure S7:** Relationships between Fox TF binding sites and mES TF binding site locations. **A)** $Log_2$ enrichment of overlap between the top 2,000 peaks for each Fox TF and peaks for the listed mES TFs. Numbers and color shading reflect the $log_2$ observed/expected enrichment of overlap between binding sites. **B)** Percentage overlap between the top 2,000 peaks for each Fox TF and peaks for the listed mES TFs.

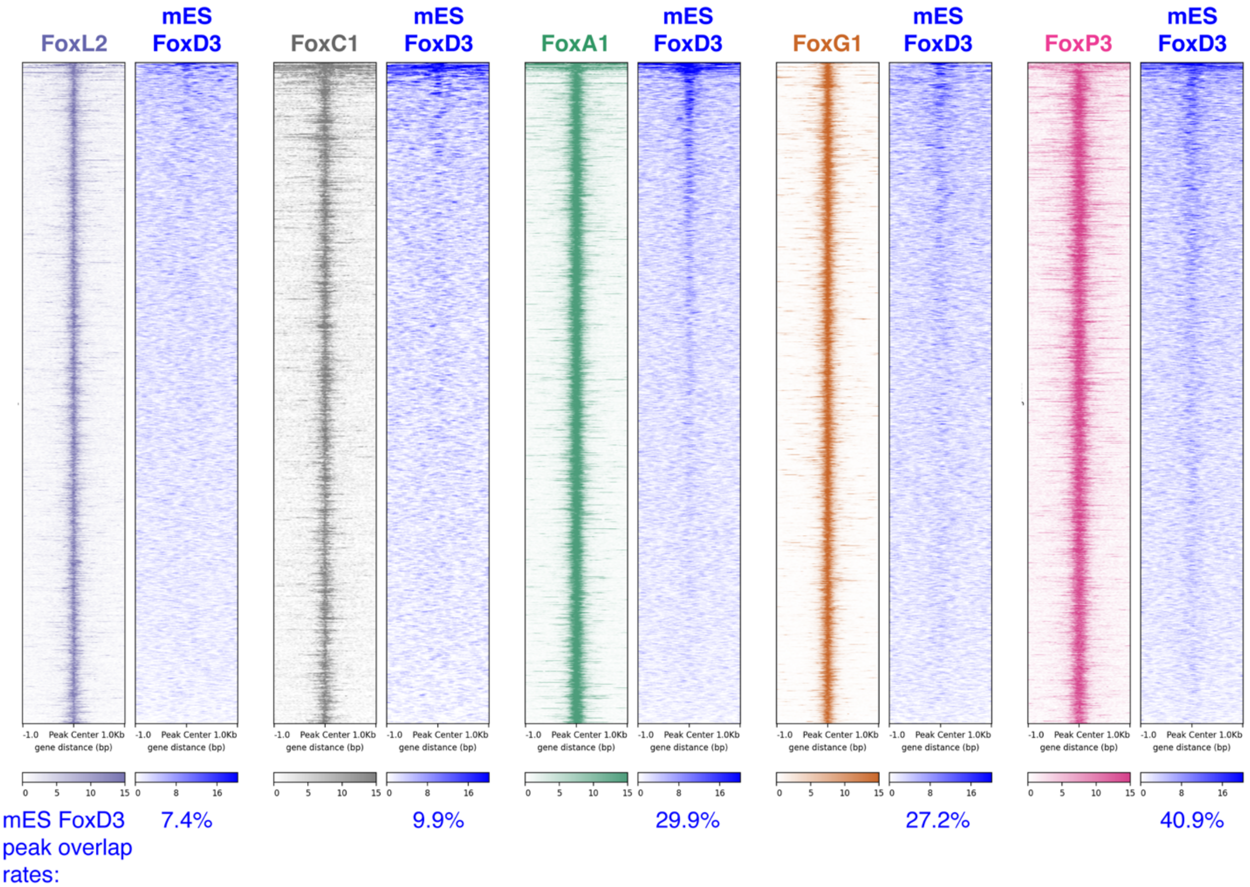

**Figure S8:** Assessing the enrichment of mES FoxD3 binding at induced Fox TF binding sites. Heatmap plots are centered on the top 2,000 ChExMix peaks for each Fox TF, and show Fox TF ChIP-exo enrichment and mES FoxD3 ChIP-seq enrichment (data from [44]). Heatmaps are sorted by decreasing FoxD3 enrichment. Percentages under each plot show the fractions of the top 2,000 induced Fox TF peaks that overlap a FoxD3 peak.

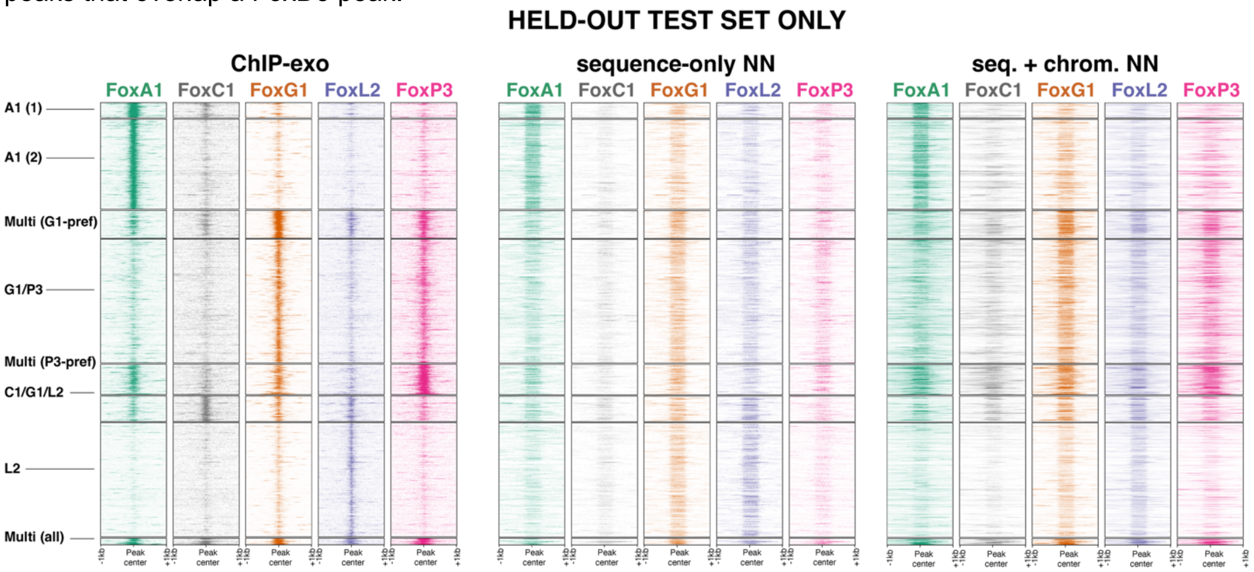

**Figure S9:** Heatmap representations of neural network predicted read counts at the clusters of Fox TF binding sites, as shown in Figure 4C, but here only displaying sites in the held-out test sets.

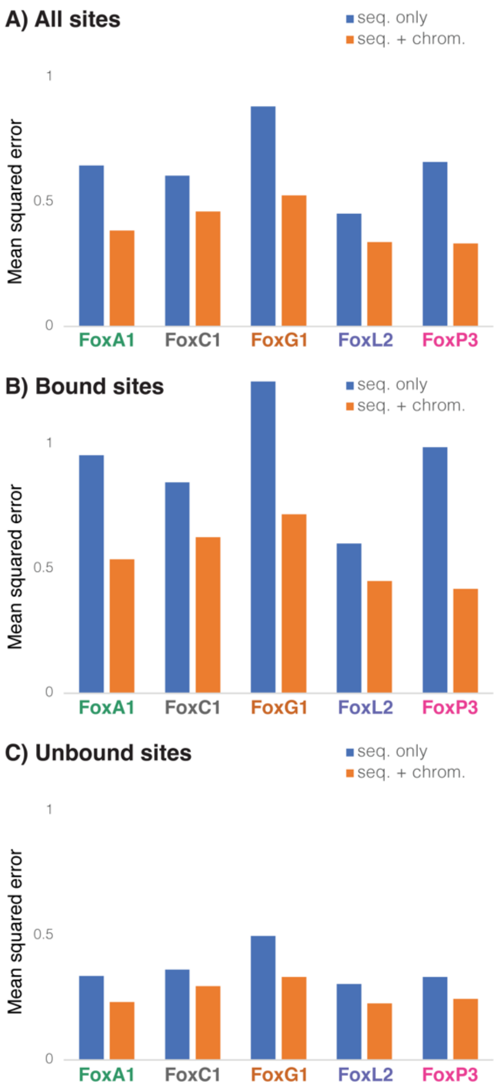

**Figure S10:** Neural network performance metrics (mean squared error) for: **A)** all held-out test set sites; **B)** bound held-out test set sites; and **C)** unbound held-out test set sites. Blue bars show the performance of networks trained only with sequence and orange bars show performance of networks trained with sequence and chromatin.

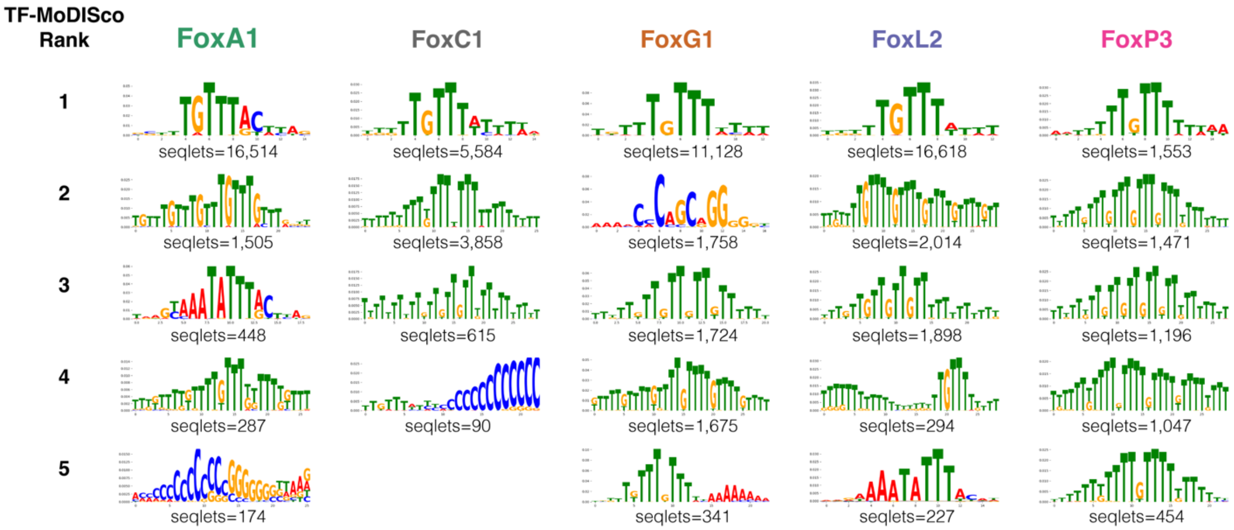

**Figure S11:** Motifs discovered by TF-MoDISco from sequence+chromatin neural network DNA sequence attribution scores for each Fox TF.

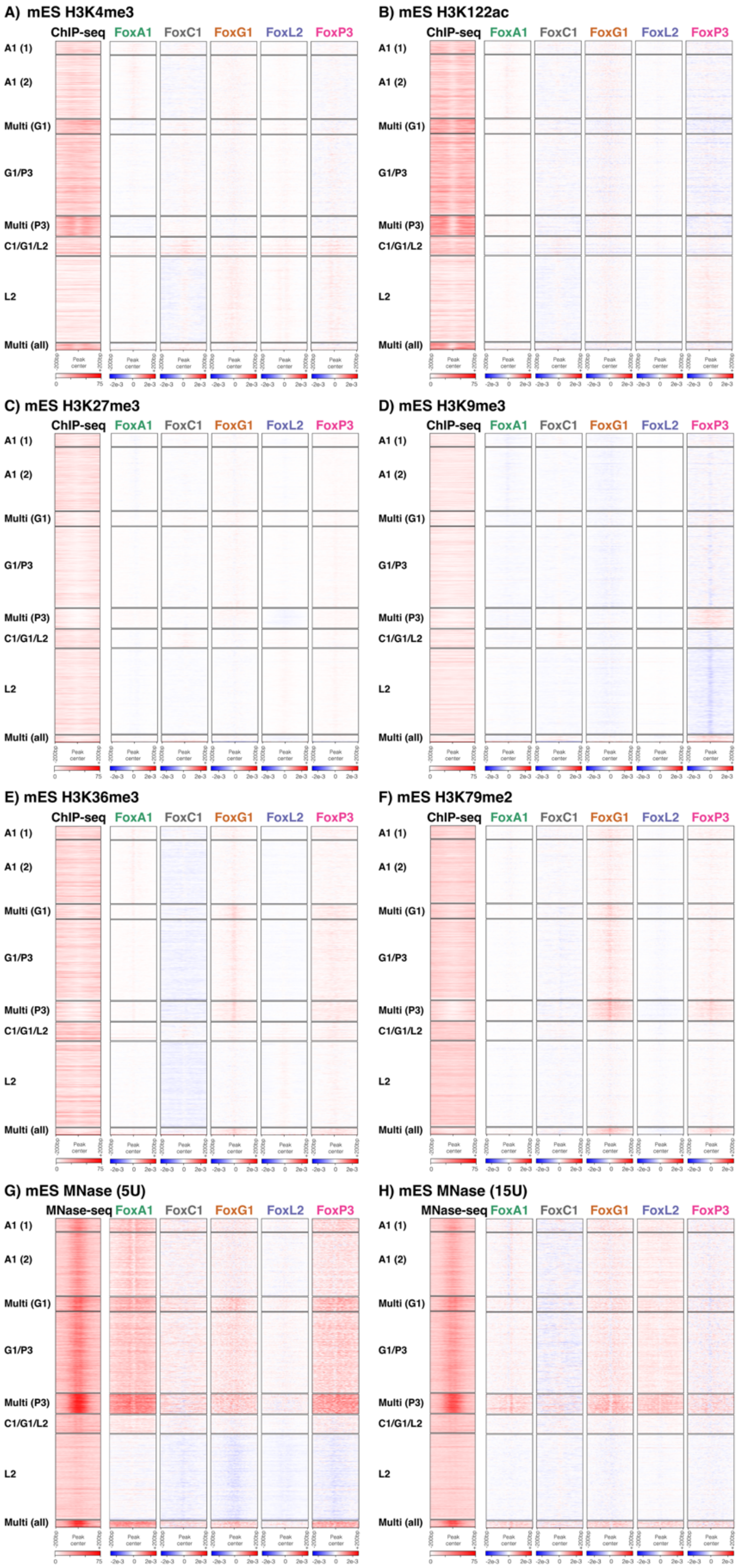

**Figure S12:** Sequence+chromatin neural network chromatin feature attribution heatmaps for clusters of Fox TF binding sites, formatted as displayed in Figure 5. Heatmaps show attributions for: **A)** mES H3K4me3; **B)** mES H3K122ac; **C)** mES H3K27me3; **D)** mES H3K9me3; **E)** mES H3K36me3; **F)** mES H3K79me2; **G)** mES MNase-seq, low concentration; **H)** mES MNase-seq, high concentration.

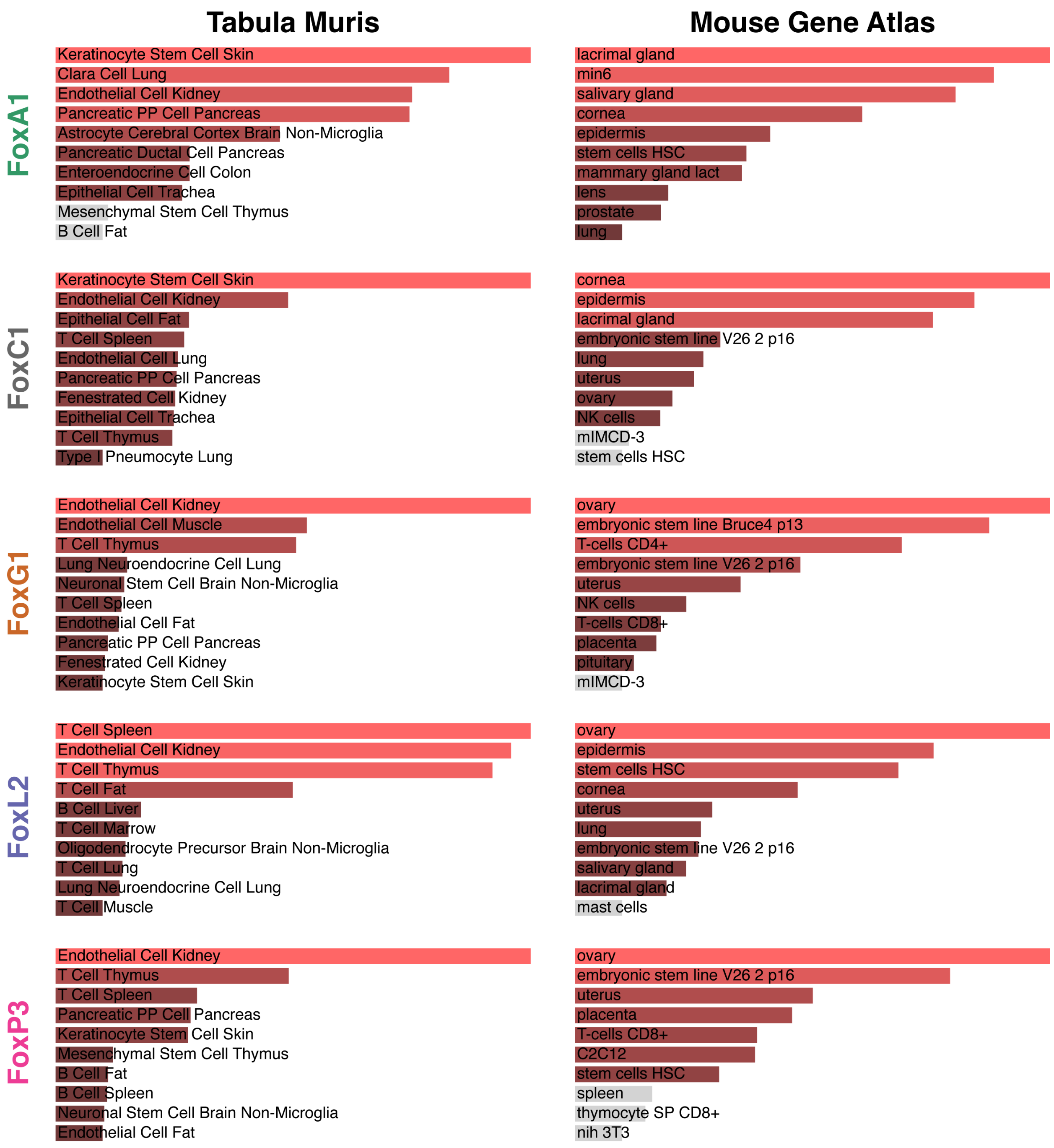

**Figure S13:** Enriched tissue-specific gene set terms from Tabula Muris and the Mouse Gene Atlas at lists of transcription factor genes that are significantly up-regulated downstream of each Fox TF.

A) iFox up-regulated genes

| Peaks | +FoxA1 | +FoxC1 | +FoxG1 | +FoxL2 | +FoxP3 |
|---|---|---|---|---|---|
| FoxA1 | 3.4 | 2.1 | 1.6 | 1.9 | 2.1 |
| FoxC1 | 2.3 | 3.2 | 1.6 | 1.8 | 1.5 |
| FoxG1 | 2.5 | 2.4 | 1.8 | 2.5 | 2.6 |
| FoxL2 | 2.0 | 2.0 | 1.6 | 2.6 | 2.1 |
| FoxP3 | 2.3 | 2.3 | 2.0 | 2.6 | 2.4 |

B) iFox down-regulated genes

| Peaks | +FoxA1 | +FoxC1 | +FoxG1 | +FoxL2 | +FoxP3 |
|---|---|---|---|---|---|
| FoxA1 | 2.0 | 2.2 | 3.3 | 2.7 | 2.0 |
| FoxC1 | 3.0 | 2.9 | 3.9 | 2.3 | 3.4 |
| FoxG1 | 2.2 | 2.5 | 3.9 | 2.4 | 2.5 |
| FoxL2 | 1.6 | 1.6 | 3.0 | 1.2 | 1.9 |
| FoxP3 | 2.9 | 2.1 | 3.5 | 2.1 | 2.6 |

**Figure S14:** Enrichment rates (observed/expected) of the top 2,000 peaks for each Fox TF nearby A) the top 250 up-regulated genes downstream of each Fox TF and B) the top 250 down-regulated genes downstream of each Fox TF.

### SUPPLEMENTAL TABLES

**Table S1:** ChExMix peak-finding results for ChIP-exo experiments performed at 0hrs, 6hrs, 12hrs, 24hrs, or 48hrs after Doxycycline induction of FoxA1 or Ascl1 in iFoxA1 or iAscl1 mES cell lines, respectively. The table shows the numbers of significant peaks discovered by ChExMix (q <0.05) and the estimated signal fraction in each experiment. Estimated signal fractions are calculated within ChExMix and are analogous to the "Fraction of Reads in Peaks" (FRiP) score. Experiments performed at 0hrs represent control ChIP-exo experiments in uninduced cell lines.

| Sample | ChExMix Peaks | Est. Signal Fraction |
|---|---|---|
| FoxA1 (native Ab) 00h | 15 | 0.006 |
| FoxA1 (native Ab) 06h | 86 | 0.009 |
| FoxA1 (native Ab) 12h | 8,345 | 0.032 |
| FoxA1 (native Ab) 24h | 35,134 | 0.096 |
| FoxA1 (native Ab) 48h | 41,726 | 0.080 |
| FoxA1 (FLAG Ab) 00h | 20 | 0.006 |
| FoxA1 (FLAG Ab) 06h | 17 | 0.014 |
| FoxA1 (FLAG Ab) 12h | 1,656 | 0.014 |
| FoxA1 (FLAG Ab) 24h | 15,984 | 0.062 |
| FoxA1 (FLAG Ab) 48h | 13,299 | 0.043 |
| Ascl1 (native Ab) 00h | 0 | 0.005 |
| Ascl1 (native Ab) 06h | 352 | 0.009 |
| Ascl1 (native Ab) 12h | 1,789 | 0.013 |
| Ascl1 (native Ab) 24h | 3,172 | 0.017 |
| Ascl1 (native Ab) 48h | 3,881 | 0.018 |

**Table S2:** Preexisting mES chromatin and TF binding data sources used for overlap analyses and chromHMM training.

| Dataset | Public ID | Publication |
|---|---|---|
| mES ATAC-seq | GSE80511 | PMID: 27939218 |
| mES H3K4me1 | GSE80482 | PMID: 27939218 |
| mES H3K4me2 | GSE80482 | PMID: 27939218 |
| mES H3K4me3 | GSE80482 | PMID: 27939218 |
| mES H3K27ac | GSE80482 | PMID: 27939218 |
| mES H3K27me3 | GSE80482 | PMID: 27939218 |
| mES Input (1) | GSE80482 | PMID: 27939218 |
| mES H3K36me3 | ENCSR000CGR | PMID: 25409824 |
| mES H3K9ac | ENCSR000CGP | PMID: 25409824 |
| mES H3K9me3 | ENCSR000ADM | PMID: 25409824 |
| mES Input (2) | ENCSR000ADJ | PMID: 25409824 |
| mES cMyc | GSE11431 | PMID: 18555785 |
| mES CTCF | GSE11431 | PMID: 18555785 |
| mES E2f1 | GSE11431 | PMID: 18555785 |
| mES Esrrb | GSE11431 | PMID: 18555785 |
| mES Klf4 | GSE11431 | PMID: 18555785 |
| mES Nanog | GSE11431 | PMID: 18555785 |
| mES nMyc | GSE11431 | PMID: 18555785 |
| mES Oct4 | GSE11431 | PMID: 18555785 |
| mES Sox2 | GSE11431 | PMID: 18555785 |
| mES Stat3 | GSE11431 | PMID: 18555785 |
| mES Tcfcp2l1 | GSE11431 | PMID: 18555785 |
| mES Zfx | GSE11431 | PMID: 18555785 |
| mES Input (3) | GSE11431 | PMID: 18555785 |
| mES Nr5a2 | GSE19019 | PMID: 20096661 |
| mES MafK | ENCSR000ESB | PMID: 25409824 |
| mES p300 | ENCSR000CCD | PMID: 25409824 |
| mES FoxD3 | GSE70545 | PMID: 26748758 |

**Table S3:** Preexisting mES chromatin data sources used in sequence+chromatin neural network training

| Dataset | Public ID | Publication |
|---|---|---|
| mES Input | GSE120376 | PMID: 31699133 |
| mES H3K122ac | GSE120376 | PMID: 31699133 |
| mES H3K27ac | GSE120376 | PMID: 31699133 |
| mES H3K27me3 | GSE120376 | PMID: 31699133 |
| mES H3K36me3 | GSE120376 | PMID: 31699133 |
| mES H3K4me1 | GSE120376 | PMID: 31699133 |
| mES H3K4me3 | GSE120376 | PMID: 31699133 |
| mES H3K79me2 | GSE120376 | PMID: 31699133 |
| mES H3K9me3 | GSE120376 | PMID: 31699133 |
| mES MNase_partial_5U | GSE82127 | PMID: 27889238 |
| mES MNase_partial_15U | GSE82127 | PMID: 27889238 |
| mES ATACSeq | GSE158528 | PMID: 34139016 |
